# KARS mediates intra-translational deposition of *N*^6^-acetyl-*L*-lysine in nascent proteins to contribute the acetylome in cells

**DOI:** 10.1101/2023.01.08.523135

**Authors:** Dingyuan Guo, Xiaoyan Zhang, Jie He, Weixin Yu, Runxin Zhou, Fuqiang Tong, Sibi Yin, Yu Wang, Yuwen Yang, Xin Xu, Long Wang, Mingzhu Fan, Shan Feng, Changbao Hong, Chuixiu Huang, Ke Liu, Zhuqing Ouyang, Yugang Wang

## Abstract

*N*^6^-acetyl-*L*-lysine residue is abundant in dietary protein but less is known about its potential influences on the diet-consumers. We herein report that KARS mediates intra- translational deposition of diet-derived *N*^6^-acetyl-*L*-lysine in nascent proteins to contribute the acetylome in cells. Acetylated dietary protein is a direct source of *N*^6^-acetyl-*L*-lysine that can widely and substantially contribute the acetylome in multiple organs of mice. By analyzing the co-crystal structure of Lysyl-tRNA synthetase (KARS) in complex with *N*^6^- acetyl-*L*-lysyl-AMP and pyrophosphate, together with *in vitro* biochemical assays, we learned that KARS can utilize *N*^6^-acetyl-*L*-lysine to produce *N*^6^-acetyl-*L*-lysyl-AMP and transfers the *N*^6^-acetyl-*L*-lysyl-moiety to lysine cognate tRNA to generate *N*^6^-acetyl-*L*- lysyl-tRNA, which introduces *N*^6^-acetyl-*L*-lysine into growing nascent polypeptide and intra-translationally results in protein acetylation. This undocumented protein modification mechanism is inherently different from post-translational modification (PTM) and termed as intra-translational modification (ITM). ITM can functionally mimic PTM mechanisms to deposit acetylation in histones to decondense chromatin. It can also modify PTM- inaccessible regions that are buried inside and functionally important to proteins. ITM is expected to extend the repertoire of acetylome and improve our understandings in protein modification modes in cells.

## Introduction

Lysine acetylation is a type of frequent and important protein post-translational modification (PTM) participating in cell signaling in biological system. In husbandry, lysine acetylation naturally occurs during the growth of animals and crops ^1–3^. In food industry, it is intensively added to dietary proteins via chemical reactions, aiming to improve the physicochemical and functional properties of commercialized proteins ^4–6^. As such, *N*^6^-acetyl-*L*-lysine residues are abundant in dietary proteins. However, the influence of consuming acetylated proteins on biological system remains unclear. *N*^6^-acetyl-*L*-lysine (AcK) is a theoretical product that derived from the digestion of acetylated proteins. It has been identified as a metabolite in mammals ^7–9^, but its functions are less studied. Whether and how *N*^6^-acetyl-*L*-lysine residues in dietary protein contribute to the acetylome in dietary protein-consumer remain to be answered.

In biological system, lysine acetylation is used to be regulated by acetyltransferases, which catalyze the post-translational addition of the acetyl-moiety from acetyl-CoA to the *N*^6^-amino-moiety of lysine residues, generating *N*^6^-acetyl-*L*-lysine residues in proteins to regulate protein functions ^10, 11^. To obtain and analyze purified recombinant proteins with site-specific lysine acetylation, Neumann et al established an *Escherichia coli* strain expressing modified PylRS-tRNACUA and lysyl-tRNA synthetase (KARS) pair that can accommodate *N*^6^-acetyl-*L*-lysine in the catalytic pocket^12^, which directly deposit *N*^6^-acetyl- *L*-lysine into polypeptides to produce recombinant proteins with acetylation at defined sites. It suggests the opportunity of *N*^6^-acetyl-*L*-lysine being intra-translationally introduced into protein synthesis, resulting in acetyltransferases-independent acetylation of nascent proteins. Whether this intra-translational introduction of *N*^6^-acetyl-*L*-lysine in nascent proteins naturally occurs in eukaryotic cells remains to be studied.

Curious about the potential contribution of diet-derived *N*^6^-acetyl-*L*-lysine to the acetylome in dietary protein-consumer, we firstly fed mice with deuterium-labelled acetylated protein and confirmed that acetylated dietary protein is a direct source of *N*^6^- acetyl-*L*-lysine that can widely and substantially contribute to the acetylome in organs of mice, including liver, brain, and lung. *N*^6^-acetyl-*L*-lysine in nutrient environment can be rapidly transported into cells. Direct incorporation of deuterium-labelled acetyl-*L*-lysine into^13^C6,^15^N2-*L*-lysine-labelled proteome was observed in mammalian cells. Co-crystal structure of Lysyl-tRNA synthetase in complex with *N*^6^-acetyl-*L*-lysyl-AMP and pyrophosphate, together with findings from biochemical assays, demonstrate that KARS can utilize *N*^6^-acetyl-*L*-lysine to produce *N*^6^-acetyl-*L*-lysyl-tRNA, which subsequently participates in protein synthesis and introduces *N*^6^-acetyl-*L*-lysine into growing nascent polypeptide, resulting in intra-translational protein acetylation. Different from the well-known mechanisms for protein post-translational modification (PTM), KARS-mediated deposition of acetyl-*L*-lysine in nascent polypeptides leads to protein acetylation in the middle of protein translation, we therefore termed this undocumented mechanism for protein modification as intra-translational modification (ITM). ITM can functionally mimic PTM mechanisms to deposit acetylation in histones to decondense chromatin, and can also modify proteins in folded and buried regions that are PTM-inaccessible, affecting protein functions via disturbing inter-residue interactions inside proteins.

### *N*^6^-acetyl-*L*-lysine residues in dietary protein contribute the acetylome of diet- consumer

To explore the cross-talk between the *N*^6^-acetyl-*L*-lysine residues in dietary proteins and the ones in the acetylome of dietary protein-consumer, we produced and fed the deuterium-labelled acetylated protein (*N*^6^-(acetyl*-d3*)-*L*-lysine-protein) to mice and collected blood samples to performe high-performance liquid chromatography-tandem mass spectrometry (HPLC-MS/MS) analysis (Figure S1A and S1B). Identification of *N*^6^- (acetyl*-d3*)-*L*-lysine (*d3*-AcK) in the blood samples built a direct connection between acetylated dietary protein and the metabolite of AcK in protein-consumer (Figure 1A and 1B; Figure S1C). To understand whether the diet-derived *d3*-AcK could contribute to the acetylome in organs and tissues of protein-consumer, we collected the liver, brain, and lung from the studied mice and analyzed the acetylation profiles. Besides the non-labelled acetylation sites, we identified 867, 387, and 480 deuterium-labelled acetylation sites in liver, brain, and lung tissues, respectively (Figure 1C to 1E; Table S1), revealing the contribution of diet-derived AcK to the acetylome in protein-consumer.

**Figure 1.**
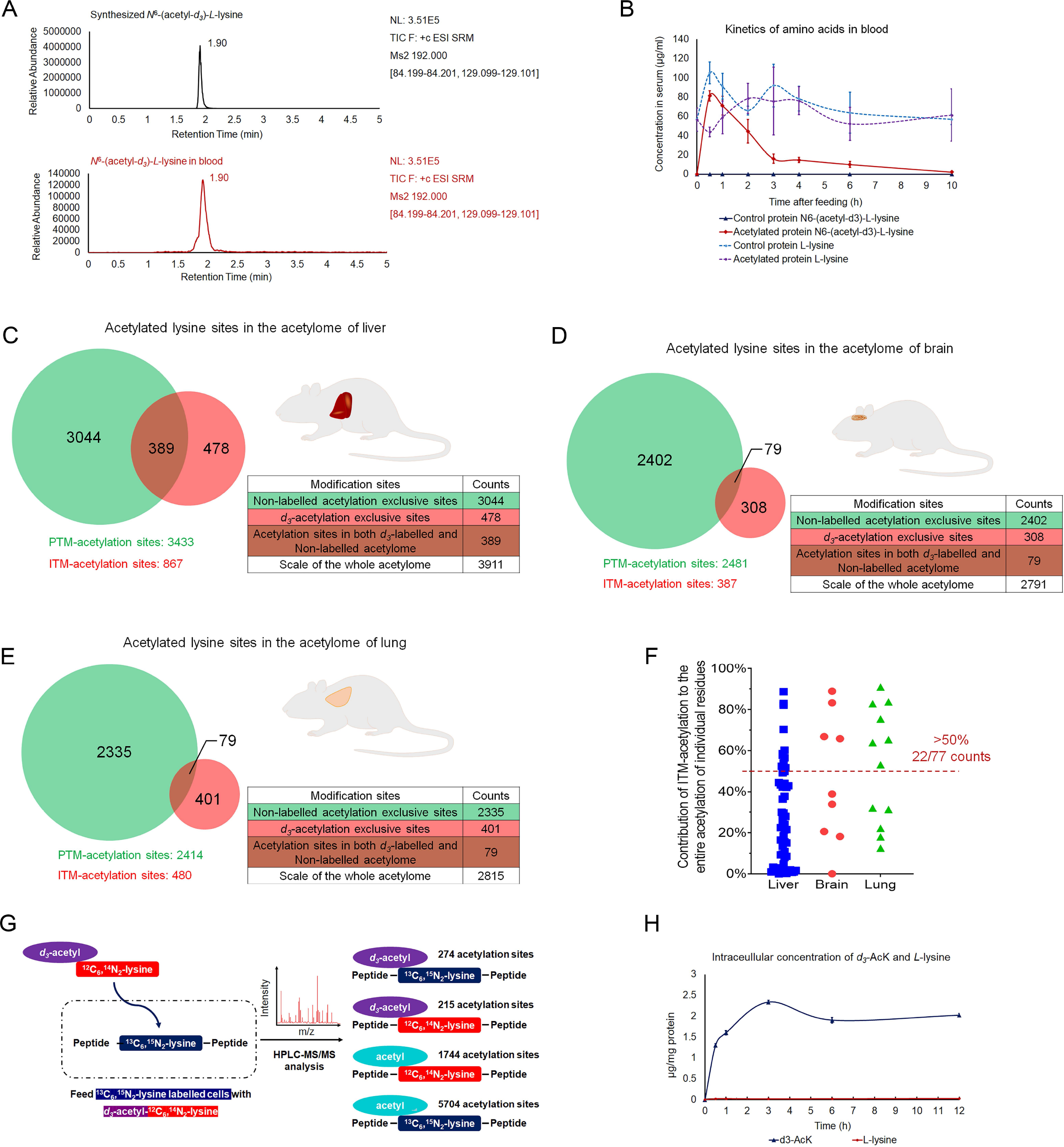
*N*^6^-acetyl-*L*-lysine residues in dietary proteins contribute the acetylome of diet-consumer. (**A**), Identification of *N*^6^-(acetyl-*d3*)-*L*-lysine in the blood of mice fed with *N*^6^-(acetyl-*d3*)- *L*-lysine-protein. *N*^6^-(acetyl-*d3*)-*L*-lysine in the blood samples and chemical synthesized *N*^6^-(acetyl-*d3*)-*L*-lysine show similar elution time in the same HPLC-MS/MS system. (**B**), The kinetics of *N*^6^-(acetyl-*d3*)-*L*-lysine and lysine in the blood of mice fed with *N*^6^- (acetyl-*d3*)-*L*-lysine-protein and control protein, respectively. Each data is presented as the means±s.d. of five mice (n=5). (**C**) to (**E**), Analyses of the acetylome of organs in the mice fed with *N*^6^-(acetyl-*d3*)-*L*- lysine-protein. The *N*^6^-(acetyl-*d3*)-modified lysine residues are ITM-acetylation sites. The *N*^6^-acetyl-modified lysine residues are PTM-acetylation sites. The ITM-acetylation sites and PTM-acetylation sites that were identified in the liver (**C**), brain (**D**), and lung (**E**) are represented, respectively. (**F**), Quantitative analysis of the contribution of ITM-acetylation to the entire acetylation of individual lysine residues. To avoid the biases inherent in the use of antibody-based enrichment methods that employed prior to the proteomic analysis ^31^, only the pairs of ITM-acetylated peptides and PTM-acetylation peptides with the same features (length, amino acid sequence, modifications and modification sites) were used for the quantitative analyses. Details for calculating the contribution of ITM-acetylation to the entire acetylation of each lysine residues are described in the part of **materials and methods**. (**G**), Identification of the *N*^6^-(acetyl-*d3*)-*L*-lysine residues in the 6C13,2N15-*L*-lysine- labelled cells. The left panel shows the scheme of *N*^6^-(acetyl-*d3*)-*L*-lysine incorporating in the 6C13,2N15-*L*-lysine-labelled proteome. The right panel shows the scheme of the HPLC-MS/MS identified peptides that are resulted from the direct incorporation of *N*^6^-(acetyl- *d3*)-*L*-lysine in 6C13,2N15-*L*-lysine-labelled proteins and the PTM-resulted protein acetylation in the 6C13,2N15-*L*-lysine-labelled cells. The number of identified acetylation positions are presented. The detail information about the identified acetylation positions is listed in the supplementary table S5. (**H**), Quantification of *N*^6^-(acetyl-*d3*)-*L*-lysine and *L*-lysine in cells. Kinetic studies of *N*^6^- (acetyl-*d3*)-*L*-lysine and *L*-lysine in ^13^C6,^15^N2-*L*-lysine-labelled cells treated with *N*^6^- (acetyl-*d3*)-*L*-lysine. The cells were cultured in the medium suppled with 1 mM ^13^C6,^15^N2- *L*-lysine. Each data is presented as the means±s.d. of three assays (n=3).

By analyzing the relative abundance of deuterium-labelled acetylation to the entire acetylation of specific lysine residues that were identified in both deuterium-labelled acetylome and non-labelled acetylome, we found about 28.6% of the acetylated lysine residues are majorly affected by deuterium-labelled acetylation (accounts more than 50% of the entire acetylation) (Figure 1F). The maximum contribution of deuterium-labelled acetylation to the entire acetylation of individual lysine residues reached 90.9% (Figure 1F; Table S2), showing the substantial contribution of diet-derived AcK to the acetylation of certain lysine residues in the acetylome of protein consumer. We next performed biological processes enrichment analyses and found that deuterium-labelled acetylation occurs on 556 proteins that are enriched in 32 biological processes in the studied liver tissue (Figure S2; Table S3), 314 proteins that are enriched in 15 biological processes in the studied brain tissue (Figure S3; Table S3), and 333 proteins that are enriched in 9 biological processes in the studied lung tissue (Figure S4; Table S3), suggesting that food-derived AcK has wide influence on the acetylome and biological processes in animal. Most of the deuterium-labelled acetylation enriched biological processes and non-labelled acetylation enriched processes were overlapped, suggesting that diet-contributed acetylome is regulated to match physiological states^10^, despite that the regulatory mechanisms might be different. These results ascertain the model that *N*^6^-acetyl-*L*-lysine bridges acetylated dietary protein and the acetylomes in dietary protein-consumer.

To understand how AcK contribute the acetylome in mammalian cells, we labelled the proteome of human liver cells with ^13^C6,^15^N2-*L*-lysine and treated the labelled cells with deuterium-labelled AcK (*N*^6^-(acetyl*-d3*)-*L*-^12^C6,^14^N2-lysine, *d3*-AcK) (Figure S1D and S1E). Proteomic analyses identified 274 *N*^6^-(acetyl*-d3*)-^13^C6,^15^N2-lysine-labelled acetylation sites (Figure 1G; Table S4), suggesting the deuterium-labelled acetyl-moiety might be released from the *d3*-AcK and recycled to form *d3*-acetyl-CoA, which latterly contribute the acetylome via acetyltransferase-dependent mechanisms ^13^. We also identified 1744 *N*^6^- acetyl-^12^C6,^14^N2-lysine acetylation sites (Figure 1G; Table S4), suggesting that *d3*-AcK might also be a source of lysine that can participate protein synthesis and contribute to the proteome ^13^. Notably, we also identified 215 *N*^6^-(acetyl*-d3*)-^12^C6,^14^N2-lysine-labelled acetylation sites (Figure 1G; Table S4), which might be resulted from recombination of the recycled *d3*-acetyl-moiety and lysine residue might serve as the explanation, or *d3*-AcK can be directly incorporated into the proteome in cells. Release of lysine and *d3*-acetyl-moiety from *d3*-AcK is the prerequisite for the recombination of the recycled *d3*-acetyl-moiety and lysine residue in the studied proteome. However, feeding *N*^6^-(acetyl*-d3*)-*L*-^12^C6,^14^N2-lysine to the ^13^C6,^15^N2-*L*-lysine-labelled cells did not significantly contribute the intracellular *L*- ^12^C6,^14^N2-lysine within 12 hours (Figure 1H). Of noted, immunoblotting analysis revealed that the acetylation level of Lys-34 and Lys-46/47 of histone H2B, Lys-118/119 and Lys- 124 of histone H2A, which were identified as *N*^6^-(acetyl*-d3*)-^12^C6,^14^N2-lysine-labelled acetylation sites, were upregulated by AcK within 12 hours (Figure S1F). Consistent with the report that histone hyperacetylation induces chromatin decondensation ^14^, the chromatin in cells treated with AcK for 12 hours was decondensed (Figure S1G). These results strongly suggest that the *d3*-acetyl-moiety and lysine residues in the labelled proteome are not directly generated from *d3*-AcK. Direct incorporation in proteome might be the primary mechanism underlying AcK contributing the acetylome in mammalian cells.

### *N*^6^-acetyl-*L*-lysine is an alternative substrate of KARS

Neumann et al established an *Escherichia coli* strain expressing modified lysyl-tRNA synthetase (KARS) that accommodates *N*^6^-acetyl-*L*-lysine (AcK) in the catalytic pocket ^12^, allowing AcK being directly deposited into nascent polypeptides to produce proteins with acetylation. This system inspired us proposing the hypothesis that KARS might mediate the direct incorporation of AcK into eukaryotic proteome. To test this hypothesis, we analyzed the structure of human wild-type KARS in complex with *L*-lysine and ATP (PDB ID: 3BJU) ^15^. Docking analysis revealed that AcK fits well in the substrate binding pocket and the predicted AcK-interacting residue profile was similar with that of lysine (Figure 2A to 2C), shedding light on the possibility of AcK being a substrate of human wild-type KARS. We next established *in vitro* enzymatic activity assays by incubated purified KARS and tRNAs with AcK. AcK-dependent ATP consumption and AMP production suggests that AcK might be an alternative substrate of KARS (Figure 2D and 2E). Docking analyses illustrated that the residues of Glu301 and Arg323 interact with both AcK and lysine in the substrate binding pocket (Figure 2C). Replacement of these residues with alanine (KARS- E301A and KARS-R323A) significantly reduced both AcK and *L*-lysine dependent ATP consumption by KARS (Figure S5A and S5B). The Thr337 residue was predicted as the one that specifically mediate KARS interacting with AcK (Figure 2C). Replacement of Thr337 with alanine (KARS-T337A) significantly reduced the AcK-dependent ATP consumption by KARS while keeping its *L*-lysine-dependent ATP consumption activity intact (Figure S5A and S5B). Steady state kinetics analyses revealed that the maximum velocity (Vmax) and Michaelis Constant (*Km*) of AcK-dependent KARS is 1.55±0.39 µM·minute^-1^ and 74.15±25.64 µM, respectively. Because KARS can catalyze the formation of *L*-lysyl-AMP that attacks the *N*^6^-amino of *L*-lysine to generate *L*-lysyl-*L*- lysine and release AMP ^16, 17^, we observed ATP consumption and AMP production by KARS in the absence of tRNAs (Figure S5C and S5D). As such, the Km and Vmax of the KARS-*L*-lysine and KARS-AcK assays cannot be precisely compared. However, we confirmed the activity of KARS utilizing AcK, tRNA, and ATP to produce AMP, which is similar with the reaction that KARS places on *L*-lysine.

**Figure 2.**
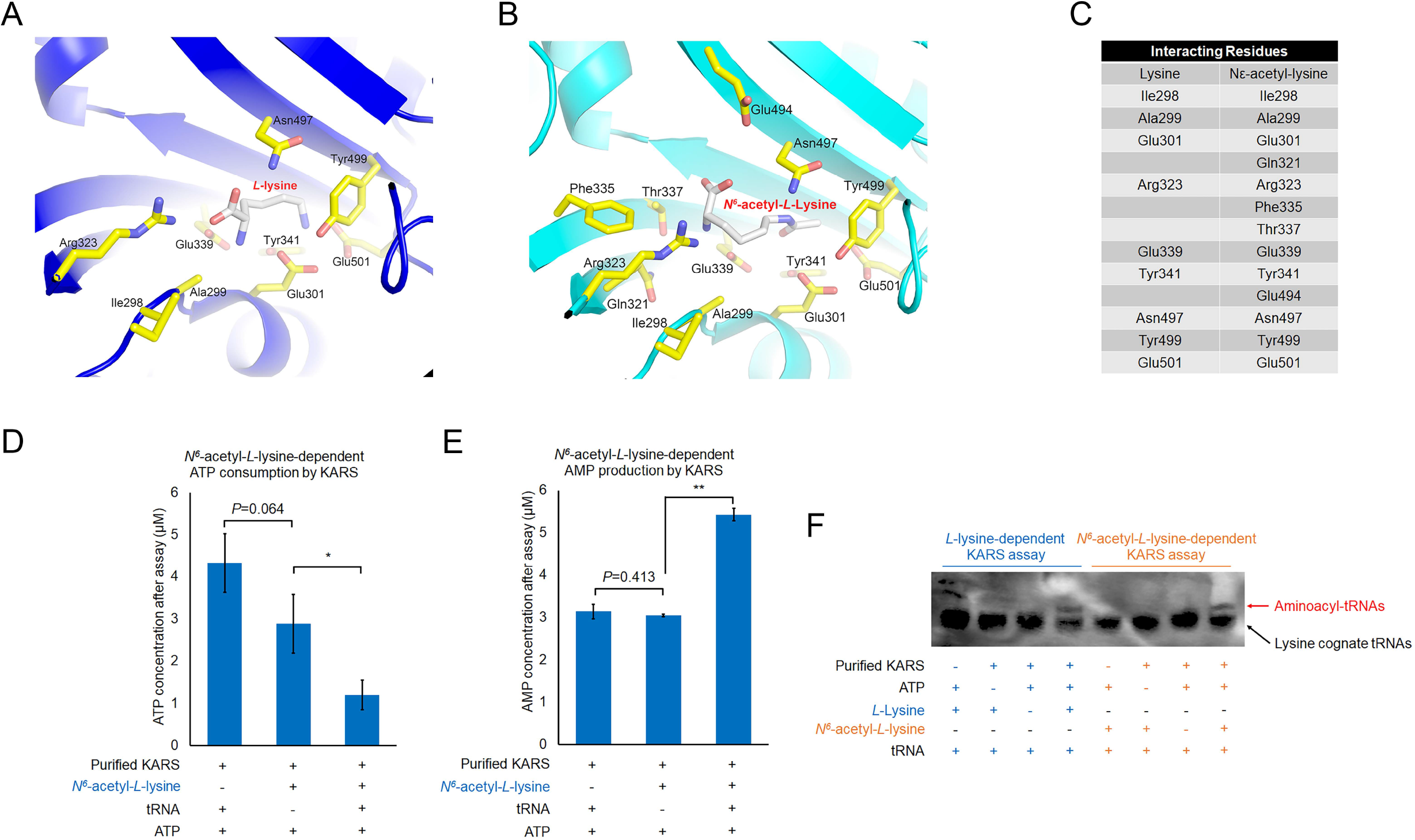
*N*^6^-acetyl-*L*-lysine is an alternative substrate of KARS. (A) to (**C**), *N*^6^-acetyl-*L*-lysine can be accommodated in the substrate binding pocket of KARS. The substrate-binding pocket of human KARS structure (PDB code: 3BJU) (**A**) was referred to perform the docking structure of KARS in complex with *N^6^*-acetyl-*L*-lysine (AcK) (**B**). *L*-lysine and *N^6^*-acetyl-*L*-lysine are shown as sticks. The residues that interact with *L*-lysine and *N^6^*-acetyl-*L*-lysine are shown as sticks, labeled in the zoomed-in view of the substrate-binding pocket of human KARS, and listed (**C**), respectively. (**D**) and (**E**), KARS can utilize *N*^6^-acetyl-*L*-lysine to catalyze the conversion of ATP into AMP. Purified KARS was incubated with ATP, *N*^6^-acetyl-*L*-lysine, and tRNAs for 1 hour. The ATP consumption (**D**) and AMP production (**E**) in the assays were measured and quantitatively analyzed. Two-sided *t*-test analyses were conducted. The data is presented as the means±s.d. of three independent experiments (n=3). \**P*<0.05. (**F**), KARS catalyzes the formation of aminoacyl-tRNAs by using lysine and *N*^6^-acetyl-*L*- lysine, respectively. Northern blotting analysis of lysine cognate tRNAs by using fluorescence-labelled single strand DNA probe that specifically recognizes lysine cognate tRNAs. The free lysine cognate tRNA and aminoacyl-tRNAs are indicated with black and red arrows, respectively. Representative images of triplicate experiments are shown.

*L*-lysyl-tRNA is the product of *L*-lysine-dependent KARS catalysis. To test if KARS could catalyze similar reaction on AcK, we performed northern blotting assays following KARS-mediated *in vitro* reactions. The mobility shift of lysine cognate tRNAs in both the AcK and *L*-lysine dependent assays strongly suggest the activity of KARS producing *N*^6^- acetyl-*L*-lysyl-tRNAs by utilizing AcK (Figure 2F; Figure S5E), supporting the possibility that KARS mediates the incorporation of AcK in nascent proteins via producing *N*^6^-acetyl- *L*-lysyl-tRNAs.

### Molecular basis of KARS utilizing *N*^6^-acetyl-*L*-lysine as substrate

To understand how KARS utilizing AcK as substrate, we explored the molecular basis of KARS-catalyzed production of *N*^6^-acetyl-*L*-lysyl-tRNA by crystalizing KARS (KARS- apo) and the complex obtained from the incubation of KARS, AcK, and ATP. Instead of the expected co-crystal of KARS in complex with AcK and ATP, we unexpectedly obtained the one representing KARS in complex with *N*^6^-acetyl-*L*-lysyl-AMP and pyrophosphate (KARS-AcK-AMP, 2.26 Å, deposited in PDB, PDB ID: 8HYQ) (Figure 3A; Table S6). Compared to the sequential mechanism that KARS generates *L*-lysyl-AMP, transfers the *L*-lysyl-moiety to tRNAs, and release *L*-lysyl-tRNAs (Figure 3B) ^18^, the co-crystal structure of KARS-AcK-AMP demonstrates that KARS can perform similar reaction on AcK and produce *N*^6^-acetyl-*L*-lysyl-AMP. The aminoacyl-tRNA that was generated from the incubation of KARS and AcK is therefore confirmed as *N*^6^-acetyl-*L*-lysyl-tRNA (Figure 2F).

**Figure 3.**
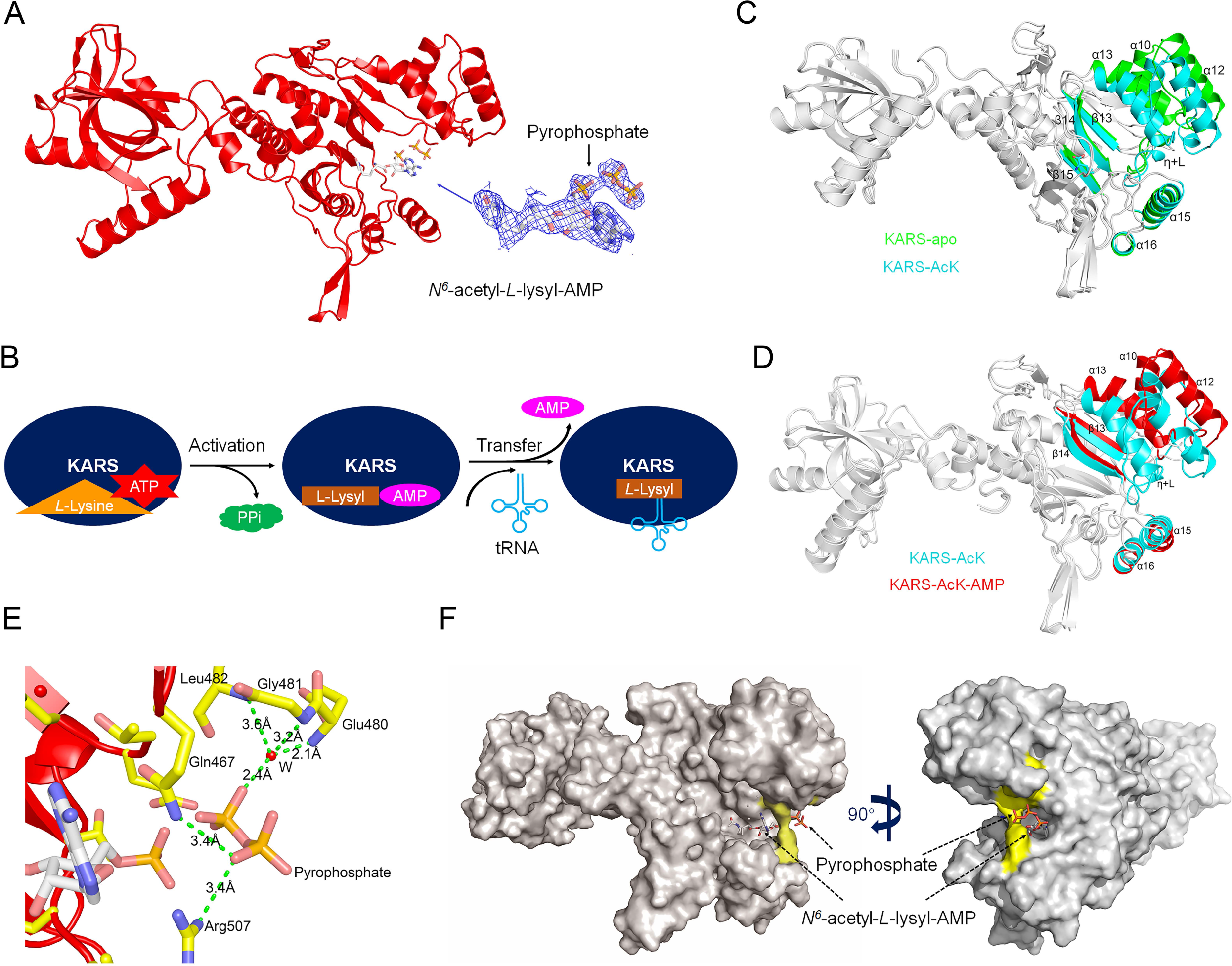
The molecular basis of KARS utilizing *N*^6^-acetyl-*L*-lysine as substrate. (**A**), Co-crystal structure of KARS in complex with *N*^6^-acetyl-*L*-lysyl-AMP and pyrophosphate. Cartoon drawing of KARS-AcK-AMP structure with zoomed in view of *N*^6^-acetyl-*L*-lysyl-AMP and pyrophosphate. The ligands are shown as sticks, along with their 2*Fo*-*Fc* electron density map (blue mesh) contoured at 1.0 σ. (**B**), The scheme of the molecular process of KARS catalyzing the production of *L*-lysyl- tRNA by using *L*-lysine. KARS catalyzes the specific attachment of L-lysine to its cognate tRNA in a 2-step reaction. L-lysine is first activated by ATP to form L-lysyl-AMP and pyrophosphate is a byproduct of this step. Then the L-lysyl-moiety is transferred from *L*- lysyl-AMP to lysine cognate tRNA to form L-lysyl-tRNA. (**C**), Structural difference between KARS in the states of standby and substrate-binding. Superimposition the structures of KARS-apo and KARS-AcK is performed. Only the regions showing structural difference between the KARS-apo (green) and KARS-AcK (cyan) are colored and labelled, including the helixes of α-10, α-12, α-13, α-15, and α-16, the helix η and its associated loop (η+L), and the sheets of β-13, β-14, and β-15. *N^6^*-acetyl- *L*-Lysine is shown as sticks. (**D**), Structural difference between KARS in the states of substrate-binding and products- releasing. Superimposition the structures of KARS-AcK and KARS-AcK-AMP is performed. Only the regions showing structural difference between the KARS-AcK (cyan) and KARS-AcK-AMP (red) are colored and labelled, including the helixes of α-10, α-12, α-13, α-15, and α-16, the helix η and its associated loop (η+L), and the sheets of β-13 and β-14. The compounds of *N^6^*-acetyl-*L*-Lysine, *N^6^*-acetyl-*L*-lysyl-AMP, and pyrophosphate are removed for better illustration of the protein structures. (**E**), Zoomed-in view of the pyrophosphate-binding site of KARS-AcK-AMP structure. Pyrophosphate-binding residues are shown as sticks and labeled. The ordered water molecule (W) and the values of the distance (green dash line) are also indicated. (**F**), Surface drawing of the crystal structure of KARS-AcK-AMP in two different orientations. *N^6^*-acetyl-*L*-lysyl-AMP and pyrophosphate are shown in sticks. Pyrophosphate-binding residues are colored in yellow.

The structures of KARS-apo (2.55 Å, deposited in PDB, PDB ID: 8HYR) (Figure S6A), KARS-AcK (Figure 3C), and KARS-AcK-AMP (Figure 3A) represent the states of KARS in standby, substrates binding, and products releasing, respectively. To understand the dynamicity of KARS in catalyzing the production of *N*^6^-acetyl-*L*-lysyl-tRNA, we analyzed the superimposition of KARS-apo and KARS-K and found a compacted catalytic domain while KARS accommodating *L*-lysine (Figure S6B). The compacted catalytic domain might facilitate the interaction between substrates for catalytic reaction (Figure S6B) ^19^. Similar conformational change was observed in the structure of KARS-AcK (Figure 3C), illustrating similar structural dynamics while KARS utilizing AcK as substrate. By analyzing the superimposition of KARS-AcK (substrates-binding) and KARS-AcK-AMP (products-releasing) (Figure 3D), we found the catalytic pocket of KARS becomes loosen. The helices of α-10, α-12, α-13, α-15, and α-16, as well as the sheets of β-13 and β-14, around the catalytic pocket were pushed outward to create an enlarged channel, which might allow the relocation of the AcK from the substrates binding position to the products releasing position (Figure S6C), along with the newly conjugated AMP moiety in the form as *N*^6^-acetyl-*L*-lysyl-AMP. The helix η and its associated loop (η+L) that originally cover the substrates are invisible in the KARS-AcK-AMP structure (Figure 3D), indicating flexibility of these regions so that they can be pushed aside to facilitate the release of *N*^6^- acetyl-*L*-lysyl-AMP. Interactions between pyrophosphate and the residues of Gln467 and Arg507 at the edge of the catalytic domain were observed (Figure 3E and 3F). These interactions might provide driving force for the release of pyrophosphate. Mutation of Gln467 and Arg507 (KARS-Q467A and KARS-R507A) significantly reduced the catalysis of the KARS-AcK reaction but the KARS-*L*-lysine reaction is unexpectedly intact (Figure S6D and S6E), suggesting a different pyrophosphate-releasing mechanism is used by KARS-K reaction ^20^. A water molecule neighboring the pyrophosphate was observed (Figure 3E). It might provide additional driving forces for pyrophosphate releasing. From substrates-binding to products-releasing, the structural dynamicity majorly occurs in the aminoacylation catalytic domain while keeping the tRNA binding domain intact (Figure S6F), suggesting limited influences of KARS-catalyzed *N*^6^-acetyl-*L*-lysyl-AMP production on its interaction with tRNAs. As such, generation of *N*^6^-acetyl-*L*-lysyl-AMP is the determinant step to produce *N*^6^-acetyl-*L*-lysyl-tRNA. These structural analyses, together with the enzymatic activity studies, lead to the conclusion that KARS can naturally utilize *N*^6^-acetyl-*L*-lysine to generate *N*^6^-acetyl-*L*-lysyl-tRNA.

### KARS-produced *N*^6^-acetyl-*L*-lysyl-tRNA results in deposition of *N*^6^-acetyl-*L*-lysine in nascent proteins

KARS-catalyzed *L*-lysyl-tRNA is an intermediary introducing *L*-lysine into growing nascent polypeptides in protein synthesis. *N*^6^-acetyl-*L*-lysyl-tRNA is likely to have similar role to introduce *N*^6^-acetyl-*L*-lysine into growing polypeptides and generate nascent proteins bearing acetylated-lysine residues. This hypothesis was supported by the finding that addition of AcK in a mammalian cell-free protein expression system dose-dependently upregulate the acetylation of synthesized protein (Figure 4A). Together with the results described above, we conclude that KRAS can utilize *N*^6^-acetyl-*L*-lysine to generate *N*^6^- acetyl-*L*-lysyl-tRNA, which introduces *N*^6^-acetyl-*L*-lysine into growing polypeptides in protein synthesis and generates nascent proteins with acetylation.

**Figure 4.**
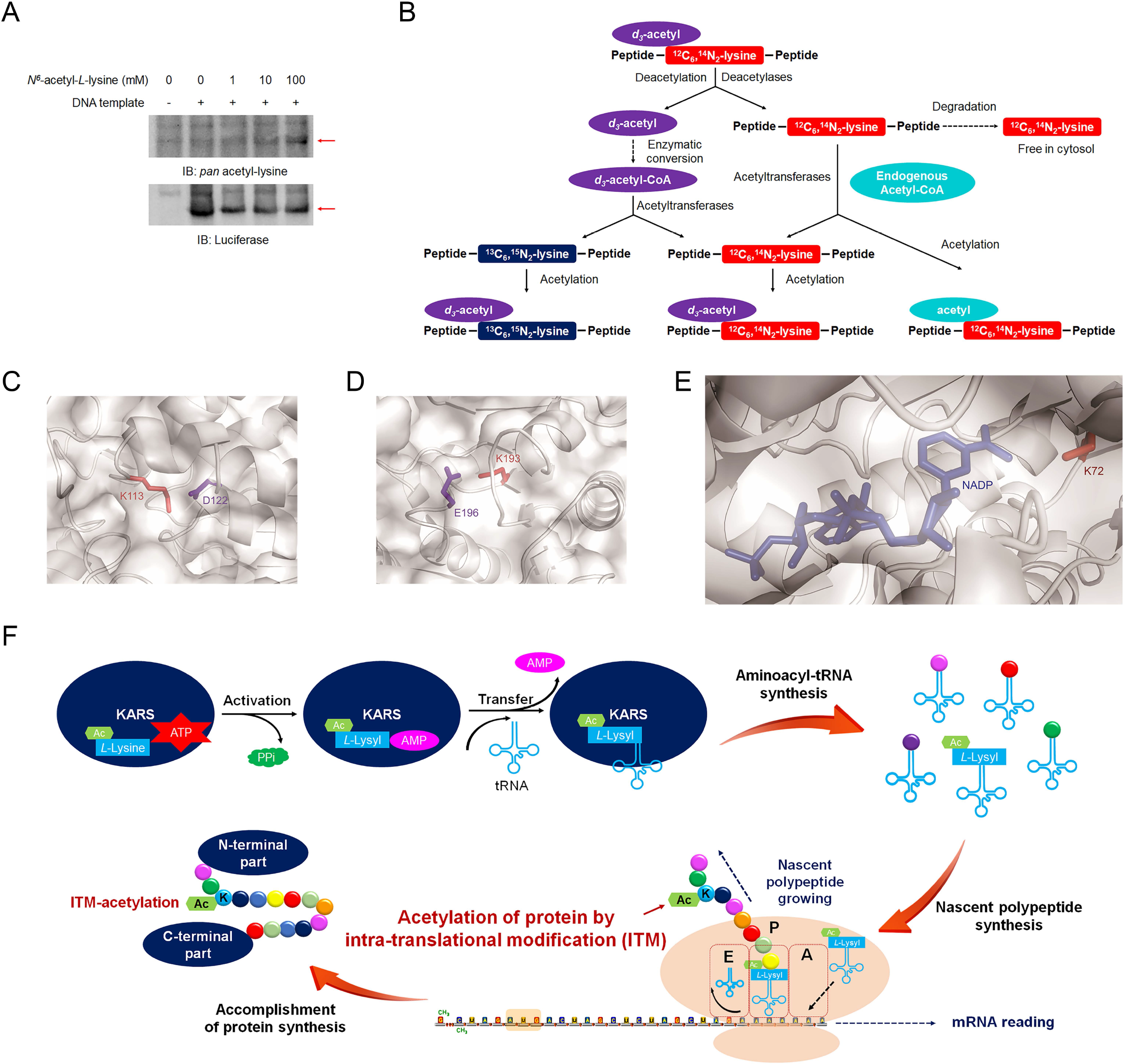
*N*^6^-acetyl-*L*-lysyl-tRNA results in deposition of *N*^6^-acetyl-*L*-lysine in nascent proteins. (**A**), *N*^6^-acetyl-*L*-lysine results in acetylation of the protein that is synthesized by cell-free protein expression system. The influence of *N*^6^-acetyl-*L*-lysine on protein synthesis was studied by the gradient addition of *N*^6^-acetyl-*L*-lysine in a mammalian cell-free protein expression system. The synthesized protein and its acetylation level were analyzed by performing immunoblotting assays with the indicated antibodies. Representative images of triplicate experiments are shown. (**B**), Scheme of the pathways for the stable isotope labelled and non-labelled acetyl-moiety and L-lysine exchange in the studied acetylomes. (C) and (**D**), Zoomed in and transparent view of mouse ALDH1A1 structure (PDB code: 7YOB). The acetylated Lys113 residue (**C**) and Lys193 residue (**D**) that localize in the buried regions of ALDH1A1 are illustrated. The acetylation affected lysine residues are shown as sticks and highlighted in red. The negative charged Asp122 and Glu196 that interact with Lys113 and Lys193 are shown as sticks and highlighted in blue, respectively. (**E**), Zoomed in and transparent view of mouse IDH1 structure (PDB code: 5YZH). The acetylated Lys72 residue that localizes in the buried regions of IDH1 is shown as sticks and highlighted in red. The substrate NADP is highlighted in blue. (**F**), Scheme of the KARS-mediated intra-translational deposition of acetyl-lysine in nascent protein results in protein acetylation.

The direct incorporation of AcK into the proteome can serve as the explanation for the acetylated peptide profiles identified in the labelled mammalian cells (Figure 1G). Because of the direct deposition of AcK in nascent proteins, *N*^6^-(acetyl*-d3*)-*L*-^12^C6,^14^N2-lysine sites were identified in the *d3*-AcK-treated ^13^C6,^15^N2-*L*-lysine-labelled cells (Figure 1G and 4B). Because the deuterium-labelled acetyl-moiety can be removed by deacetylase and leave the *L*-^12^C6,^14^N2-lysine residues that can be re-acetylated by PTM mechanisms ^13, 21^, the *N*^6^- acetyl-^12^C6,^14^N2-lysine sites were identified (Figure 1G and 4B). The removed acetyl-moiety could be recycled as *d3*-acetyl-CoA that can be utilized in acetylation of other lysine residues ^22, 23^, so that *N*^6^-(acetyl*-d3*)-^13^C6,^15^N2-lysine sites were identified (Figure 1G and 4B). Degradation of both *N*^6^-(acetyl*-d3*)-*L*-^12^C6,^14^N2-lysine and *N*^6^-acetyl-^12^C6,^14^N2-lysine bearing proteins can release the nonlabelled-lysine, so that the delayed identification of lysine in the *d3*-AcK treated ^13^C6,^15^N2-*L*-lysine-labelled cells (Figure 1H and 4B).

Another line of evidence supporting the direct incorporation of AcK in proteome is that notable amount of deuterium-labelled acetylation sites localizing inside the buried regions of proteins were identified in the studied acetylomes (Table S5), such as retinal dehydrogenase (ALDH1A1, protein ID: P24549, Lys113 and Lys193) and cytoplasmic isocitrate dehydrogenase (IDH1, protein ID: O88844, Lys72) (Figure 4C to 4E). These acetylated sites are inaccessible to the acetyltransferases, which are required for protein acetylation via PTM mechanisms. Because direct incorporation of AcK during the growth of nascent polypeptide occurs prior to the folding of proteins, it makes acetylation of these buried lysine residues possible. These findings collectively lead to the conclusion that direct incorporation of AcK in nascent protein naturally occurs in mammal, resulting in acetyltransferases-independent protein acetylation.

Taken together, the present study demonstrates that *N*^6^-acetyl-*L*-lysine directly bridges the *N*^6^-acetyl-*L*-lysine residues in dietary protein and the acetylomes in dietary protein- consumer. Direct incorporation of AcK into nascent proteins is an undocumented and important mechanism that AcK contribute the acetylome in cells. KARS can utilize *N*^6^- acetyl-*L*-lysine to produce *N*^6^-acetyl-*L*-lysyl-tRNA, which introduces AcK into nascent polypeptide synthesis, resulting in protein acetylation prior to the accomplishment of protein synthesis (Figure 4F). Compared with the well-known PTM mechanisms that occur after protein synthesis ^24^, this undocumented mechanism leads to protein acetylation in the middle of protein translation, it is therefore termed as intra-translational modification (ITM). ITM-acetylation naturally and widely occurs in multiple organs. It provides us with new understandings in the mechanisms regulating protein modification and extends the repertoire of acetylome in cells.

## Discussion

While diet-derived *N*^6^-acetyl-*L*-lysine can directly contribute to the acetylome in mammalian system, daily consumption of acetylated dietary protein might not significantly alternate the acetylome and physiologies of the diet-consumer. In the present study, we noted that AcK treatment showed limited influence on the level of pan-acetylation in cells (Figure S7A). This negative response might be resulted from the monitor of deacetylases that remove “unfavorable” acetylation to ensure the acetylomes coupling with the physiological states of cells ^25, 26^. For the same reason, most of the deuterium-labelled acetylation enriched biological processes and non-labelled acetylation enriched processes were overlapped (Figure S2-S4). Dysfunction of the deacetylases-dependent surveillance unleashed AcK-contributed acetylation in cells (Figure S7B). Deacetylases inhibitors are important candidates for diseases treatment ^27^. The cross-talk between acetylated dietary protein and deacetylases inhibitors in patients might be alerted. Moreover, physiological influences of daily consumption of acetylated dietary protein are expected to be watched.

However, some ITM-acetylated lysine residues might escape from deacetylases’ surveillance. Because ITM co-translationally occurs and installs acetylation prior to the accomplishment of protein synthesis, any lysine residues in the proteome, including the one localized in the buried and PTM-inaccessible regions, can theoretically be modified by ITM-acetylation. These ITM-acetylated PTM-inaccessible regions are also beyond the surveillance of deacetylases. They might disturb the inter-residue interactions inside proteins and influence protein functions and stabilities ^28^. Some affected proteins might be mis-folded and degraded due to protein quality-control mechanism^29^. Other portion of the affected proteins might stay in cells with alternated activities. For instance, Lys113 and Lys193 are buried inside the structure of ALDH1A1 (Figure 4C and 4D) ^30^. Acetylation can neutralize the positive charge of Lys113 and Lys193 that might disrupt their interactions with Asp122 and Glu196 (Figure 4C and 4D), respectively. We expressed and purified the proteins of ALDH1A1 wild-type and acetylation mimic mutants (ALDH1A1- K113Q and ALDH1A1-K193Q) (Figure S8A). Acetylation-mimic mutation of Lys113 and Lys193 significantly disrupted the enzymatic activity of ALDH1A1 (Figure S8B). Thermofluor shift assays revealed lower Tm of the ALDH1A1-K193Q mutant than that of wild-type (Figure S8C), suggesting ITM-acetylation of Lys193 disrupts the stability of ALDH1A1 protein. Acetylation of Lys72 of IDH1 is another illustrative example. We expressed and purified the proteins of IDH1 wild-type and acetylation mimic mutant IDH1- K72Q (Figure S8D). Steady kinetic studies revealed that IDH1-K72Q has higher Km (95.22±54.67 μM) toward NADP than that of wild-type IDH1 (16.45±5.97 μM) (Figure S8E). The maximum velocity (Vmax) of utilizing NADP to produce NADPH by IDH1- K72Q (2.43±0.45 μM·minute^-1^) is significantly lower than its wild-type counterpart (11.52±1.53 μM·minute^-1^) (Figure S8F and S8G). We therefore conclude that the acetylation-mimic mutation could disrupt the binding of IDH1 to NADP and reduce the enzymatic activity. Moreover, IDH1-K72Q mutant showed lower thermo-stability than IDH1wild-type in thermofluor shift assays (Figure S8H), suggesting ITM-acetylation on the buried Lys72 residue could reduce the stability of IDH1. As such, it is worthy asking the accumulative effect of acetylation in the buried regions of proteins on cells.

While ITM-acetylation is demonstrated and provides us with a new mode for protein modification that might be parallel with PTM-mechanism, there are at least three facets to be refined to sculpture the outline of ITM-acetylation. First, studies showing the influence of ITM-acetylation on physiologies in details are wanted. Second, besides buried regions, lysine residues on the surface of proteins can also be acetylated by ITM mechanism, so that ITM-acetylation can cross-talk with PTM-acetylation. However, the mechanisms underlying the cross-talk remain to be studied. Third, protein quality control system might involve in the surveillance of ITM-acetylation on “unwanted” sites in nascent proteins, so that only small number of ITM-acetylation sites were identified in the PTM-cold regions. The surveillance mechanism remains to be explored. In addition to acetylation, other types of protein modification are also likely to be installed by ITM mechanism. Relevant reports are expected to diversify the field of protein intra-translational modification in future.

## Supporting information

PDB validation report-8HYQ

PDB validation report-8HYR

## Acknowledgements

We sincerely thank Dr. Hongtao Yu at the Westlake University for his scientific input in this work. We thank Mingzhu Fan and Shan Feng at the Mass Spectrometry & Metabolomics Core Facility of Westlake University for proteomic analyses. We thank the staff from BL18U1 beamline of National Facility for Protein Science in Shanghai (NFPS) at Shanghai synchrotron Radiation Facility, for assistance during data collection. This work was supported by National Natural Science Foundation of China (91957110 and 31970577 to Y.W, 31870751 to Z.O), the National Key Research and Development Program of China (2018YFE0204500 to Z.O).

## Author Contributions

Y.W. and Z.O conceived and designed the study and wrote the manuscript. D.G., X.Z., W.Y., Y.W., F.T., S.Y., and S.L. performed the biochemical experiments to identify the KARS activities *in vivo* and *in vitro*. J.H., C.H., and C.H., performed HPLC-MS/MS to quantitatively analyze *N*^6^-acetyl-*L*-lysine and *L*-lysine in cells and blood samples. R.Z. and K.L. performed bioinformatics analyses. X.X. and L.W performed the chemical synthesis of *N*^6^-(acetyl*-d3*)-*L*-lysine. M.F. and S.F. performed proteomic analyses. X.Z. and Z.O. performed structural analyses.

## Author Information

The authors declare no competing financial interests.

## Materials and Methods

### Materials

Primary antibodies for immunoblotting: β-actin (mouse, Cat#AC004, lot#3500100006, Abclonal, Wuhan, China), Acetylation-lysine (rabbit, Cat#9441S, lot#15, Cell Signaling Technology, MA, USA), Luciferase (rabbit, Cat#A18259, lot#3501634001, Abclonal, Wuhan, China). The SILAC DMEM medium (Cat#88425) were purchased from Thermo fisher (Shanghai, China) and prepared according to the manufacturer’s guideline. Fluorescent probe for detecting lysine cognate tRNA probe (5’Cy5.5-ATT AAG AGT CAT ACG CG) was synthesized and purchased from Sangon Biotech (Shanghai, China). Nucleic acid gel stain Solargreen (Cat#G5570) was purchased from Solarbio (Beijing, China). Northern blot hybridization solution (Cat#AM8663) and nylon membrane (Cat#AM10100) were purchased from Thermo Fisher Scientific (Shanghai, China). Diethyl Pyrocarbonate (Cat#V900882) and tRNA from brewer′s yeast (Cat#10109525001) were purchased from Sigma-Aldrich (Shanghai, China). *N^6^*-acetyl-*L*-lysine (Cat#N159376) were purchased from Aladdin (Aladdin, Shanghai, China). Six-week-old female ICR and C57BL/J were purchased from Hubei Bainte Biological Technology Co., Ltd.

### Cell culture and stable isotope labeling

Human hepatocyte LO2 cells were kindly provided by Dr. Weidong Xie at Tsinghua University. The cells were cultured in Dulbecco’s modified Eagle’s medium (Cat#SH30285.01, Cytiva Life Sciences, MA, USA) supplemented with 10% fetal bovine serum (Cat#086-150, WISENTINC, Quebec, Canada) and 1% penicillin and streptomycin (Cat#BL505A, biosharp, Beijing, China). The cultured cells were kept at 37℃ in the cell culture incubator (CLM-170B-8-NF, ESCO, Singapore) with 5% CO2.

To label the proteome of cultured cells, cells were labeled and maintained in SILAC DMEM medium that was supplemented with 1mM ^13^C6 N2-*L*-lysine (Cat#608041, Sigma- Aldrich, Shanghai, China), 10% dialyzed fetal bovine serum, and 1% penicillin and streptomycin. After ten cell doublings, the labelling efficiency was more than 99.5%.

### Synthesis of *N^6^*-(acetyl-*d3*)-*L*-lysine

To obtain *N^6^*-(acetyl-*d3*)-*L*-lysine, *L*-Lysine was first dissolved in NaHCO3 aqueous solution. To the given solution, CuSO4·5H2O in water and NaHCO3 were added. The mixture was cooled to 0 ℃ by ice bath and acetic anhydride-*d6* (Cat#175641, Sigma- Aldrich, Shanghai, China) was added dropwise. After stirred at room temperature for 12 h, a pale blue suspension was formed. After filtration, the pale blue solid was suspended in water, then 8-hydroxyquinoline was added. The suspension was stirred for 10 h, and the color changed from blue to yellow. After filtration, the filtrate was washed with ethyl acetate. The aqueous phase was freeze-dried to give the with solid product (yield: 28%). The products were identified by the ^1^H NMR spectrum.

### Acetylation of soy proteins

The acetylation of soy proteins was performed according to the protocol that was previously reported ^32^. Briefly, soy protein (Cat#9010-10-0, Shanghai Macklin Biochemical Co., Ltd, Shanghai, China) was homogenized with 0.1M carbonate buffer, pH 8.3, for 3 h at room temperature. Acetic anhydride-*d6* (Cat#175641, Sigma-Aldrich, St. Louis, USA) was gradually added to the soy protein isolate solution (anhydride/protein = 0.6-1g/g) and carried out for 1h. The solution pH was adjusted with 2.5M NaOH to be in the range of 8.0-8.5. After the pH has stabilized, the solution was kept at room temperature for 2h, and then dialyzed with double-distilled water at 4°C for 48h to remove excess anhydride. The dialyzed protein was freeze-dried and stored at -20°C.

### Acetyl-peptide enrichment and identification

The ^13^C6 N2-*L*-lysine-labelled LO2 cells (>6×10) were treated with 10mM *N* - (acetyl-*d3*)-*L*-lysine for 48h. Then the cells were collected and washed three times with ice- cold PBS. Mice were daily fed with 0.1g *N^6^*-(acetyl*-d3*)-*L*-lysine-protein for 5 days. Organs and tissues were collected and washed three times with ice-cold PBS and stored at -80°C for later analysis.

Collected cells or tissue samples (>0.2g each) were lysed and homogenized with urea lysis buffer (20 mM HEPES pH 8.0, 9 M urea, 1×protease and phosphatase inhibitor cocktail, 1×deacetylase Inhibitor Cocktail). Lysates were sonicated (60-Watt output with 6 bursts of 10 sec each, ice-bath for 30 sec between each burst) to shear genomic DNA. The protein concentrations were determined using BCA protein assay kit. 20 mg protein from each sample was processed for further procduces.

Acetylated peptides were enriched following the PTMScan kit (Cell Signaling Technologies, Danvers, MA, USA) protocol with some modifications. Briefly, proteins were reduced with 5 mM dithiothreital at room temperature for 60 min followed by alkylation with 20 mM iodoacetamide in dark for 30 minutes at room temperature. Proteins were diluted 4-fold with 20 mM HEPES pH 8.0 and then digested with trypsin at the enzyme-to-substrate ratio of 1:75 overnight at room temperature with gentle mixing. Peptides from the digestion reactions were desalted using Oasis HLB cartridge and lyophilized. To enrich acetyl-peptides, incubating the sample with PTMScan® acetyl- lysine motif immunoaffinity beads overnight at 4℃. The beads were washed twice with immunoaffinity purification buffer and three times with mass spectrometry grade water.

Then the enriched acetyl-peptides were eluted with 100 ul of 0.15% TFA and lyophilized.

Identification of acetyl-peptides by LC-MS/MS. The enriched acetyl-peptides were analyzed with the Thermo EASY-nLC1200 integrated nano-HPLC system that was directly interfaced with the Thermo Q Exactive HF-X mass spectrometer. Peptides in the chromatographic system were eluted with solvent A (0.1% formic acid) and solvent B (80% acetonitrile and 0.1% formic acid) at a flow rate of 0.300 µL/min. Each sample was analyzed nine times (6.5μL of the sample per run). The mass spectrometer was operated in the data-dependent acquisition mode using the Xcalibur 4.1 software. There is a single full- scan mass spectrum in the Orbitrap (400-1800 m/z, 60,000 resolution) that was followed by 20 data-dependent MS/MS scans at 30% normalized collision energy. Each mass spectrum was analyzed using the Thermo Xcalibur Qual Browser and Proteome Discoverer for the database searching.

Quantitative analysis of ITM-acetylation was only performed by using the pairs of ITM-acetylated peptides and PTM-acetylation peptides with the same features (length, amino acid sequence, modifications and modification sites). The contribution of ITM- acetylation to the entire acetylation of each lysine residues was calculated by using the equation 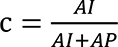, where c presents contribution of ITM-acetylation to the entire acetylation of each lysine residues, *AI* presents the abundance of the ITM-acetylated peptide that calculated in the MS system, AP presents the abundance of the PTM-acetylated peptide that has the same profile with that of ITM-acetylated peptide.

### Quantitative analysis of *N*^6^-(acetyl-*d3*)-*L*-lysine in serum

Blood samples were collected from mice fed with 0.05g *N*^6^-(acetyl*-d3*)-*L*-lysine-protein or control proteins (without acetylation treatment). The blood samples were collected from mouse’s retro-orbital sinus at the indicated time-points. The serum was isolated by centrifuging the blood at 3000rpm for 15min. 50μl serum was obtained at each time-point from each mouse. The metabolites were extracted twice with 80% methanol in water. Two extracts from the same sample were combined and dried by Vacufuge Concentrator System (Eppendorf 5301, Eppendorf, Germany).

The amino acids in mice serum were quantitatively analyzed using LC-MS/MS. 50μl of methanol/water (1:4, v/v) was applied for reconstitution. The final reconstitution samples (injection volume: 2μl) were injected into a 100×4.6mm 3.5 µm high-performance liquid chromatography column (XBridgeTM Amide, Waters, USA). The mobile phase A was consisted of 5 mM ammonium acetate in 0.1% formic acid solution, and mobile phase B was methanol organic phases. Data acquisition was performed using an Ultimate 3000 HPLC system interfaced with a TSQ Quantum Access MAX triple quadrupole Mass Spectrometry (Thermo Fisher Scientific, Waltham, MA, USA).

### Quantitative analysis of *N*^6^-(acetyl-*d3*)-*L*-lysine in LO2 cell

To analyze the intracellular concentration of *N*^6^-(acetyl-*d3*)-*L*-lysine, the cultured cells were treated with 1mM *N*^6^-(acetyl-*d3*)-*L*-lysine and collected at the indicated time-points. The collected cells were washed three times with ice-cold PBS. The metabolites were extracted with 80% methanol in water and collected following 15,000 rpm×20 minutes centrifuge. The metabolites were included in the supernatant that was dried by Vacufuge Concentrator System (Eppendorf 5301, Eppendorf, Germany). The final reconstitution samples were injected into a 100×4.6mm 3.5 µm high-performance liquid chromatography column (XBridgeTM Amide, Waters, USA). The mobile phase consisted of ammonium acetate and formic acid in both aqueous and organic phases. Data acquisition was performed using a UHPLC–LTQ/Orbitrap MS (Thermo Fisher Scientific, Waltham, MA, USA) via an Ion Trap Orbitrap positive ion mode.

### Plasmids, protein expression and purification

The KARS70–580 gene was cloned into pET-20b vector with C-terminal 6×His tag and overexpressed in *Escherichia coli* BL-21 strain. Protein expression was induced with 0.2 mM isopropyl β-D-1-thiogalactopyranoside (IPTG) at 16℃ for 20 h. Then the bacteria were harvested and resuspended in lysis buffer (20 mM Tris-HCl pH 8.0, 350 mM NaCl, 20mM imidazole, 1 mM PMSF, 1 mM benzamidine and 10 mM β-Me). The cells were lysed by “JNBIO” Low Temperature Ultra-High-Pressure Continuous Flow Cell Disrupter (JN-Mini Pro, JNBIO, Guangzhou, China) and the soluble supernatant was collected by centrifugation at 25000 g for 1 h at 4℃. The supernatant was passed through a gravity column with 2 mL Ni Focurose 6FF IMAC (HZ1003-5, HUIYAN Bio, Wuhan, China) that had previously been equilibrated with lysis buffer. Protein was then purified using a Resource^TM^ Q 6ml column (17-1179-01, GE Healthcare, Uppsala, Sweden) and a Superdex 200 Increase 10/300 column (28-9909-44, GE Healthcare, Uppsala, Sweden) equilibrated with 25 mM HEPES pH 7.5, 150 mM NaCl and 5 mM DTT. The peak fractions containing the protein were pooled and concentrated in centrifugal concentrators (CLVG01230, Merck Millipore, Cork, Ireland) to a final concentration of 30 mg/ml for crystallization. Plasmids expressing KARS-R323A, KARS-T337A, KARS-Q467A and KARS-R507A were made by site-directed mutagenesis.

The mouse IDH1 and ALDH1A1 gene were cloned into a pColdI expression vector with *N*-terminal hexahistidine tag separately and overexpressed in *Escherichia coli* T7

Express cell line (BC206-01, Biomed, Beijing, China). Protein expression was induced with 0.2 mM isopropyl β-D-1-thiogalactopyranoside at 16℃ for 20 h. Then the bacteria were harvested and protein was purified with Ni Focurose 6FF IMAC (HZ1003-5, HUIYAN Bio, Wuhan, China) and eluted with elution buffer 2 (20 mM Tris-HCl, pH 7.5, 500 mM NaCl, 300mM imidazole, pH8.0, 10% glycerol, 5 mM β-Me). Then the protein was concentrated with centrifugal concentrators (UFC901096, Merck Millipore, Cork, Ireland), and loaded onto a Superose^TM^ 6 Increase 10/300 GL gel-filtration column (29- 0915-96, GE Healthcare, Uppsala, Sweden) equilibrated with 20 mM HEPES, pH 7.5, 200 mM NaCl, 2mM DTT, 10% glycerol. The purity of the fractions containing the protein was checked by SDS-PAGE analysis. Acetylation-mimic mutants of K72Q, K113Q and K193Q were made by site-directed mutagenesis.

### Recombinant protein activity assays

KARS activity assays were performed according to the protocol that was previously described ^33^. Briefly, the reaction mixture containing 30 mM Tris-Cl pH 7.5, 150 mM NaCl, 30 mM KCl, 40 mM MgCl2, 1 mM DTT, 10 μM ATP, 3 mg/ml tRNA and 10 mM lysine or *N^6^*-acetyl-*L*-lysine were started by adding 20 μg KARS (WT/Q467A/R507A/E301A/R323A/T337A). Enzymatic activities of KARS wild-type and mutants were determined by measuring AMP production and ATP consumption with AMP-Glo Assay Kit (Cat#V5011, Promega, Madison, USA) and ATP Bioluminescent Assay Kit (Cat#FLAA, Sigma-Aldrich, Shanghai, China) or ATP Assay Kit (Cat#S0026, Beyotime, Shanghai, China), respectively.

IDH1 activity assays were performed according to the protocol that was previously reported ^34^. Briefly, the reaction mixture containing 20 mM Tris-Cl, pH 7.5, 150 mM NaCl, 10 mM MgCl2, 0.05% BSA, 2 mM β-Me, 4ng IDH1 WT/K72Q, 0.35 mM isocitrate were started by adding 0.25 μM NADP+. The IDH1-produced NADPH was measured by the diaphorase/resazurin system, which converts resazurin into resorufin (excitation 550 nm, emission 585 nm) that was quantitatively analyzed by plate reader (CLARIOstar Plus, BMG Labtech, German).

ALDH1A1 activity assays were performed according to the protocol that was previously reported ^34^. Briefly, the reaction mixture containing 50 mM Tris-Cl, pH 7.5, 200 mM NaCl, 10 mM KCl, 2 mM DTT, 200 ng ALDH1A1 WT/K113Q/K193Q, 1 mM acetaldehyde were started by adding 250 μM NAD+. The ALDH1A1-produced NADH was measured by the diaphorase/resazurin system, which converts resazurin into resorufin (excitation 550 nm, emission 585 nm) that was quantitatively analyzed by plate reader (CLARIOstar Plus, BMG Labtech, German). The concentrations of recombinant proteins were determined by Pierce™ Rapid Gold BCA Protein Assay Kit (Cat#A53225, Thermo Fisher Scientific, Rockford, USA).

### Thermofluor Shift Assay

SYPRO Orange Protein Gel Stain (Cat#S5692, Sigma-Aldrich, St. Louis, USA) were diluted 1:2500 into the assay buffer (150 mM NaCl, 200 mM Tris-Cl pH 7.5, 10 mM MgCl2, 20 mM β-Mercaptoethanol). ALDH1A1 (WT/Mutant) or IDH1 (WT/Mutant) was added with the final concentration of 0.2 μM, respectively. The 50 μl mixture were loaded into PCR 8-Strip Tubes (Cat#CP0101, GSBIO, Wuxi, China). Melting curves were obtained applying a temperature gradient from 25 to 99 °C and a heating rate of 0.5 °C/s. The fluorescence intensity was measured by a CFX Connect™ Real-Time PCR System (Cat#1855201, Bio-Rad, USA).

### Crystallization, Data Collection and Structure Determination

To obtain the co-crystal of KARS70–580 binding to *N^6^*-acetyl-*L*-lysine (AcK), protein solution of KARS70–580 was first preincubated with synthetic AcK (nKARS70–580: nAcK=1:10) and 5 mM ATP for 1 h at 4℃. The initial crystallization screening was carried out by the sitting-drop vapor-diffusion method at 16℃ using commercial screening kits. Crystals were observed after 7-8 days in a condition containing 0.06 M Morpheus Divalents, 0.1 M Morpheus Buffer system 1, pH 6.5 and 30% Morpheus Precipitant Mix 2. Single crystals were obtained by the seeding method. Crystals were harvested and subsequently soaked in mother liquor supplemented with 5 mM AcK and 5 mM ATP for approximately 11 hours. The crystals were cryoprotected with the reservoir solution supplemented with 20% (v/v) glycerol and then flash-cooled in liquid nitrogen. The space group of the crystal is P212121, with cell dimensions of a = 130 Å, b = 90 Å, and c = 107 Å. There are two molecules in the asymmetric unit with a 53% solvent content. KARS70– 580 apo form crystals were obtained in a similar condition.

X-ray diffraction data were collected from a single crystal on beamline BL17U1 and BL18U1 of National Facility for Protein Science in Shanghai (NFPS) synchrotron Radiation Facility and processed with XDS and AIMLESS^35^. The structure of KARS70–580- AcK complex was determined and refined using the phenix suite program^36^. The structure of human LysRS (PDB entry 3bju) was processed by the Sculptor program. Two ensembles, one containing residue 300-430 and the other comprising the rest part in chain A of 3bju, were used for MR search. Model building was performed using Coot^37^. Model refinement was carried out with Phenix-Refine^38^. Ramachandran plot and the quality of the structure were evaluated with MolProbity ^39^. The final models were deposited in the protein Data

Bank with accession code 7VWQ (KARS-apo) and 7VWR (KARS-AcK). PyMOL was used to present the structure. Refinement statistics are summarized in Table S6.

### Structure docking analysis

Molecular docking of *N*^6^-acetyl-*L*-lysine with KARS70–580 was performed using CB- Dock (http://cao.labshare.cn/cb-dock/), a web server used for predicting ligand-protein interaction blindly with a popular docking program ^40^. The result with the highest docking scores was used for structure analysis.

### MNase assay

Chromatin decondensation was tested as previously described (reference #13 in the manuscript). Nuclei extractions were performed by using the Nuclei PURE Prep kit (Cat#NUC201, Sigma-Aldrich, Shanghai, China). For MNase assay, 5μl nuclei were digested with micrococcal nuclease (Cat#M0247S, NEB, MA, USA) at 37℃ for gradient time. The digestion was stopped by adding 0.5M EDTA (pH 8.0). To analyze the digested genomic DNA, proteinase K(NEB) and RNase A (Solarbio) were added in the digestion products which were incubated at 37℃ for 1hour to remove protein and RNA. Genomic DNA were purified using phenol–chloroform extraction and then analyzed by 1.5% agarose gel.

### Analysis of the surface accessibility of ITM-acetylation exclusive sites

By using the protein ID in the ITM-exclusive acetylome, we found the link of the structure in the UniProt database (https://www.uniprot.org/), then obtained the PDB format structure files of the candidate proteins from the PDB database (https://www.rcsb.org/). The spatial position of the candidate lysine sites in the candidate proteins were analyzed by using Swiss-PdbViewer 4.1.0 software. If the accessibility of the lysine residues to the surface is less than 30%, it can be termed as buried inside, otherwise it was termed as outside. If the structure is not available in PDB, a homolog protein structure with high identity will be analyzed, or we analyze the burying status of the candidate lysine residues by using alphafold predicted structure (https://alphafold.ebi.ac.uk/).

### Immunoblotting

Proteins were extracted from cultured cell using 0.5% SDS lysis buffer containing protease, deacetylase and phosphatase inhibitor cocktail. Immunoblotting analyses with indicated antibodies were performed as described previously ^41^.

### Cell Free Protein Expression Coupled with *N*^6^-acetyl-*L*-lysine

The cell free protein expression assays were performed by using the TnT® Quick Coupled Transcription/Translation System (Cat#L1171, Promega, Madison, USA), according to the manufacturer’s guideline with minor modification. Briefly, the DNA template for luciferase expression is the positive control DNA template that is provided in the kit. Gradient titrations of *N*^6^-acetyl-*L*-lysine were added in the assays. The incubation condition for protein synthesis is 30℃ for 2hr. The protein expression level of luciferase is detected by immunoblotting assay with the antibody against luciferase (rabbit, Cat#A18259, lot# 3501634001, Abclonal, Wuhan, China). The acetylation level of the synthesized luciferase was measured by immunoblotting assay with the antibody against pan-acetylation lysine.

### Analysis of *L*-lysyl-tRNA and *N*^6^-acetyl-*L*-lysyl-tRNA by northern blot hybridization

KARS-catalyzed reactions were performed by mxing 10 mM Lysine or *N^6^*-acetyl-*L*-lysine, 1 mM ATP, 3 mg/ml tRNA and 40 μg KARS that were incubated at 37℃ for 2h. The produced *L*-lysyl-tRNA and *N^6^*-acetyl-*L*-lysyl-tRNA were separated from total tRNAs with acid-urea polyacrylamide gel electrophoresis and detected with the method that was previously described ^42^. Briefly, 4 μg of total tRNA mix sample was loaded to a 12% gel. The electrophoreses were performed with100 V constant voltage at 4 °C for 8.5 hours. The tRNA was transferred onto nylon membrane by TRANS-Blot SD (Cat#170-3940, Bio-Rad, USA) at 650 mA for 30 min and the membrane was washed briefly in 5 × SSC (0.75 M sodium chloride, 0.075 M sodium citrate, adjust to pH 7.0 with 1 N HCl) and cross-linked by UV using 120mJ. The membrane was hybridized overnight with Cy5.5 labelled tRNA probe at 30℃. The signal was scanned and collected by an Odyssey CLX for fluorescence detection after membrane was washed by washing buffer (6 × SSC, 0.1% SDS) for twice.

### Pathway Enrichment

Kyoto Encyclopedia of Genes and Genomes (KEGG) pathway enrichment analysis was performed via DAVID (https://david.ncifcrf.gov/), and data was visualized using the Enrichment Map v3.3.3 ^43^ and yFiles Layout Algorithms v1.1.1 plugin of Cytoscape v3.8.244.

### Statistical Analysis

The significance of differences in the experimental data was determined using Student’s *t*-test (two-tailed) unless specifically indicated. Differences in means were considered statistically significant at *P* < 0.05 or *P* < 0.01.

---

**Figure S1.**
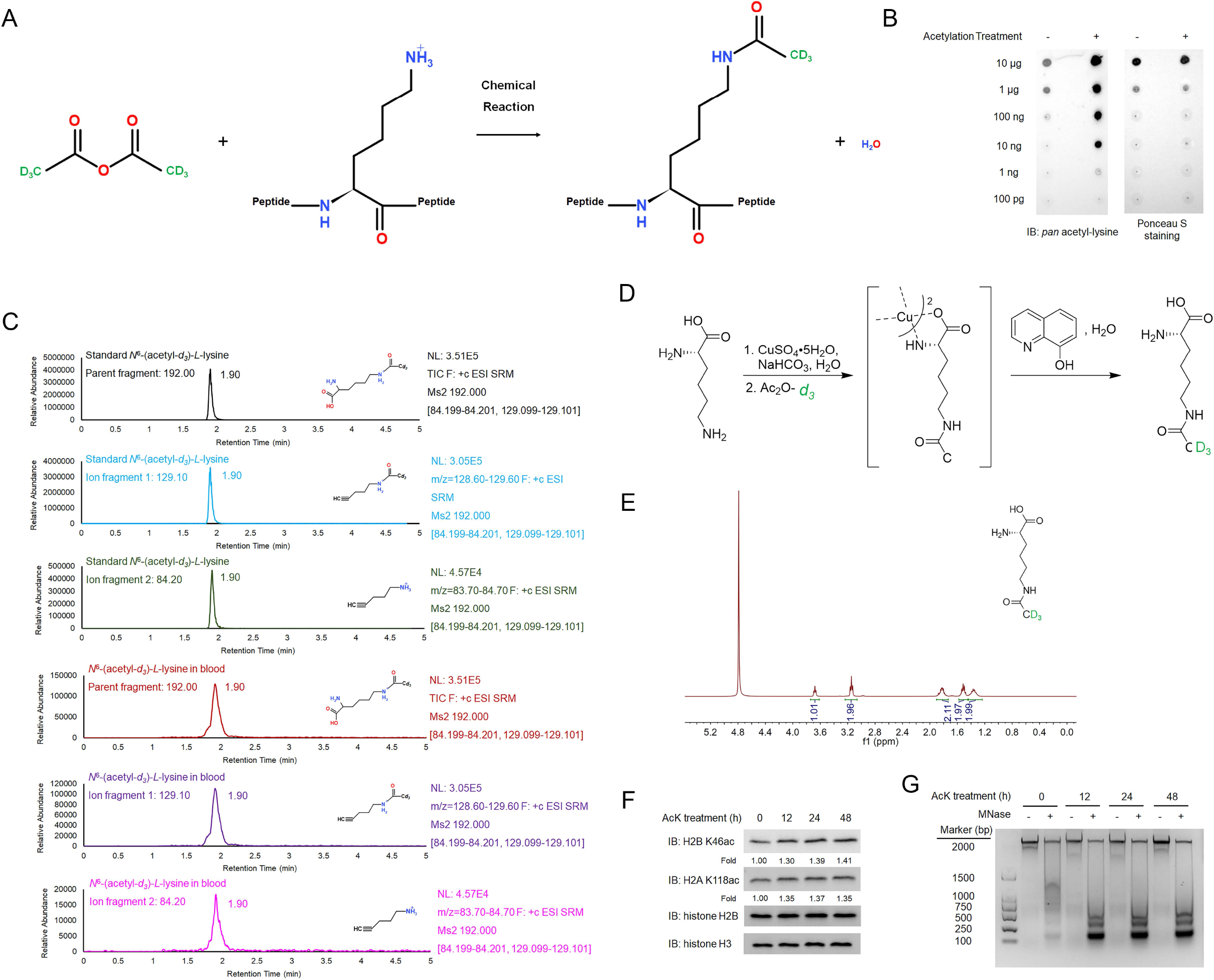
*N*^6^-acetyl-*L*-lysine residues in diet contributes the acetylome of diet- consumer. (**A**), Illustration of the workflow of synthesizing deuterium-labelled acetylated dietary protein (*N*^6^-(acetyl-*d3*)-*L*-lysine-protein). (**B**), Validation of the synthesized deuterium-labelled acetylated dietary protein by performing immunoblotting assay with the antibody against acetyl-lysine. Representative images of triplicate experiments are shown. (**C**), Identification of *N*^6^-(acetyl-*d3*)-*L*-lysine in the blood from mice fed with *N*^6^-(acetyl- *d3*)-*L*-lysine-protein. The parent ion and other two ion fragments of synthesized *N*^6^-(acetyl- *d3*)-*L*-lysine have similar elution profiles with that of *N*^6^-(acetyl-*d3*)-*L*-lysine-lysine that identified in the blood from mice fed with *N*^6^-(acetyl-*d3*)-*L*-lysine-protein. (**D**), Illustration of the workflow of synthesizing deuterium-labelled *N^6^*-acetyl-*L*-lysine (*N*^6^-(acetyl-*d3*)-*L*-lysine). (**E**), Validation of the synthesized *N*^6^-(acetyl-*d3*)-*L*-lysine. ^1^H NMR (D2O, 500 MHz, 295 K, ppm) spectrum of the synthesized *N*^6^-(acetyl-*d3*)-*L*-lysine, recorded on the Agilent spectrometer. (**F**), *N*^6^-acetyl-*L*-lysine treatment upregulates acetylation of core histones. The cultured LO2 cells were treated with 1 mM *N*^6^-acetyl-*L*-lysine for a series of time-points. The acetylation level of histone H2A K118 and histone H2B K46 in the treated cells were analyzed by immunoblotting assays with the indicated antibodies. Representative images of triplicate experiments are shown. (**G**), *N*^6^-acetyl-*L*-lysine treatment induced decondensation of chromatin in cells. The nuclei were isolated from cultured cells that had been treated with 5mM N6-acetyl-L-lysine for a series of time-points. The nuclei were then incubated with MNase, by which the genomic DNA in the decondensated chromatin regions was digested. The genomic DNA with and without MNase digestion was separated in 1.5% DNA electrophoresis agarose gel and visualized by DNA dye staining. Representative images of triplicate experiments are shown.

**Figure S2.**
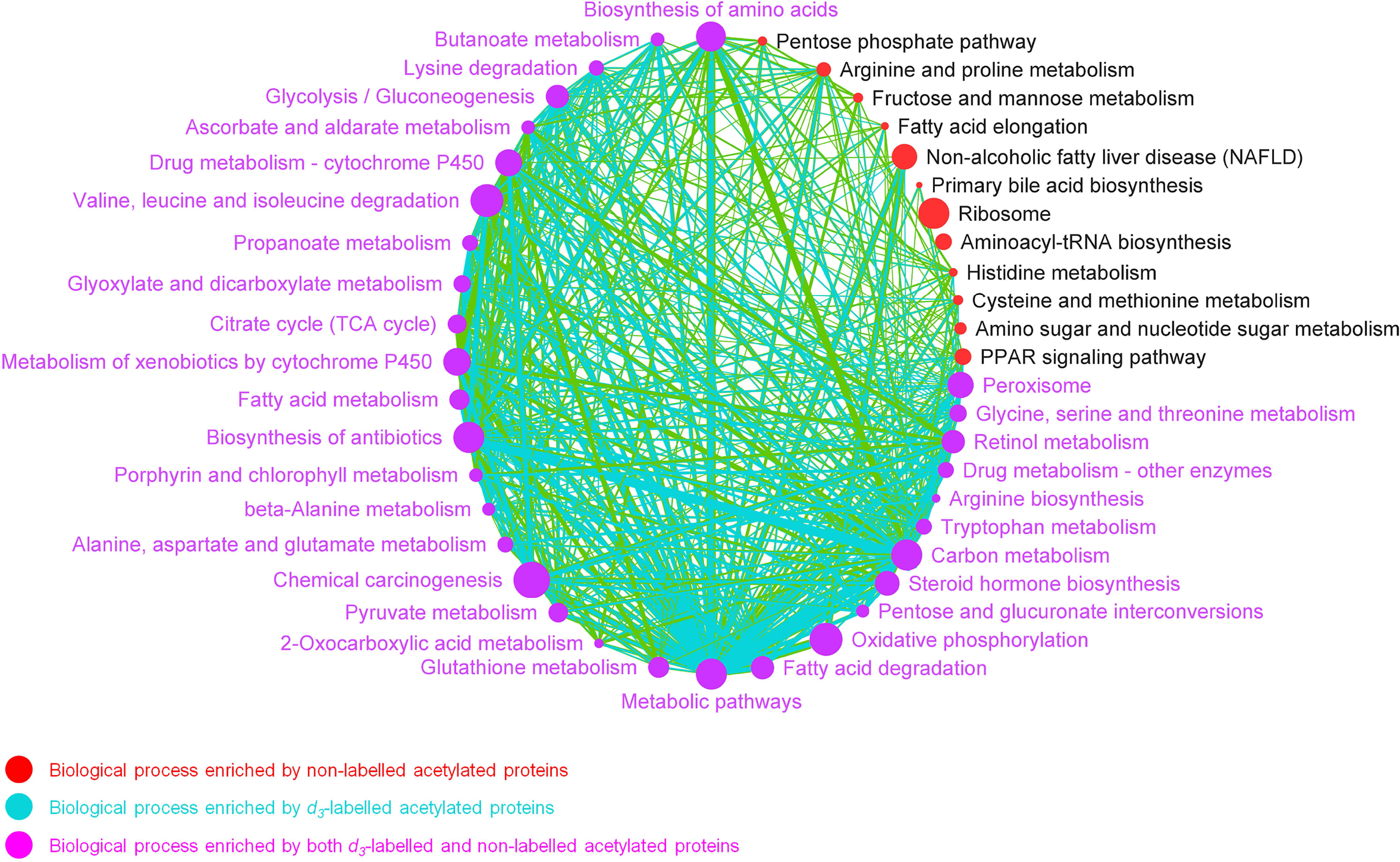
Biological processes enriched with proteins with ITM-acetylation in mice liver. Nodes stand for KEGG defined pathways (*P*<0.001 and FDR<0.001). Each node represents one biological process. The node size reflects the number of proteins in the pathway. Two nodes are connected if there were one or more proteins included in both nodes. Biological processes that are enriched by PTM-acetylated proteins (red), ITM-acetylated proteins (cyan), and proteins in both ITM-acetylome and PTM-acetylomes (plum) are highlighted in colors.

**Figure S3.**
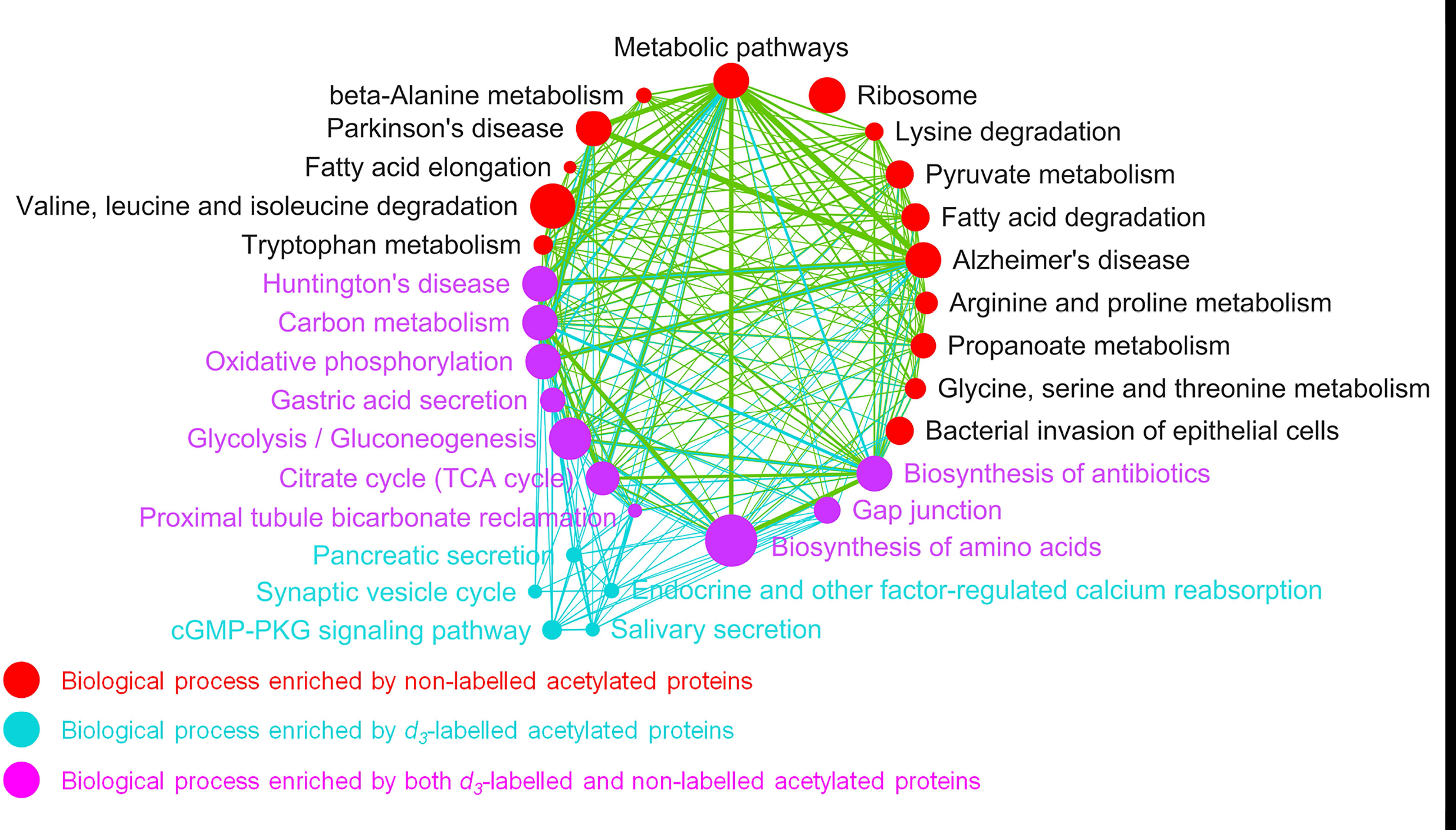
Biological processes enriched with proteins with ITM-acetylation in mice brain. Nodes stand for KEGG defined pathways (*P*<0.001 and FDR<0.001). Each node represents one biological process. The node size reflects the number of proteins in the pathway. Two nodes are connected if there were one or more proteins included in both nodes. Biological processes that are enriched by PTM-acetylated proteins (red), ITM-acetylated proteins (cyan), and proteins in both ITM-acetylome and PTM-acetylomes (plum) are highlighted in colors.

**Figure S4.**
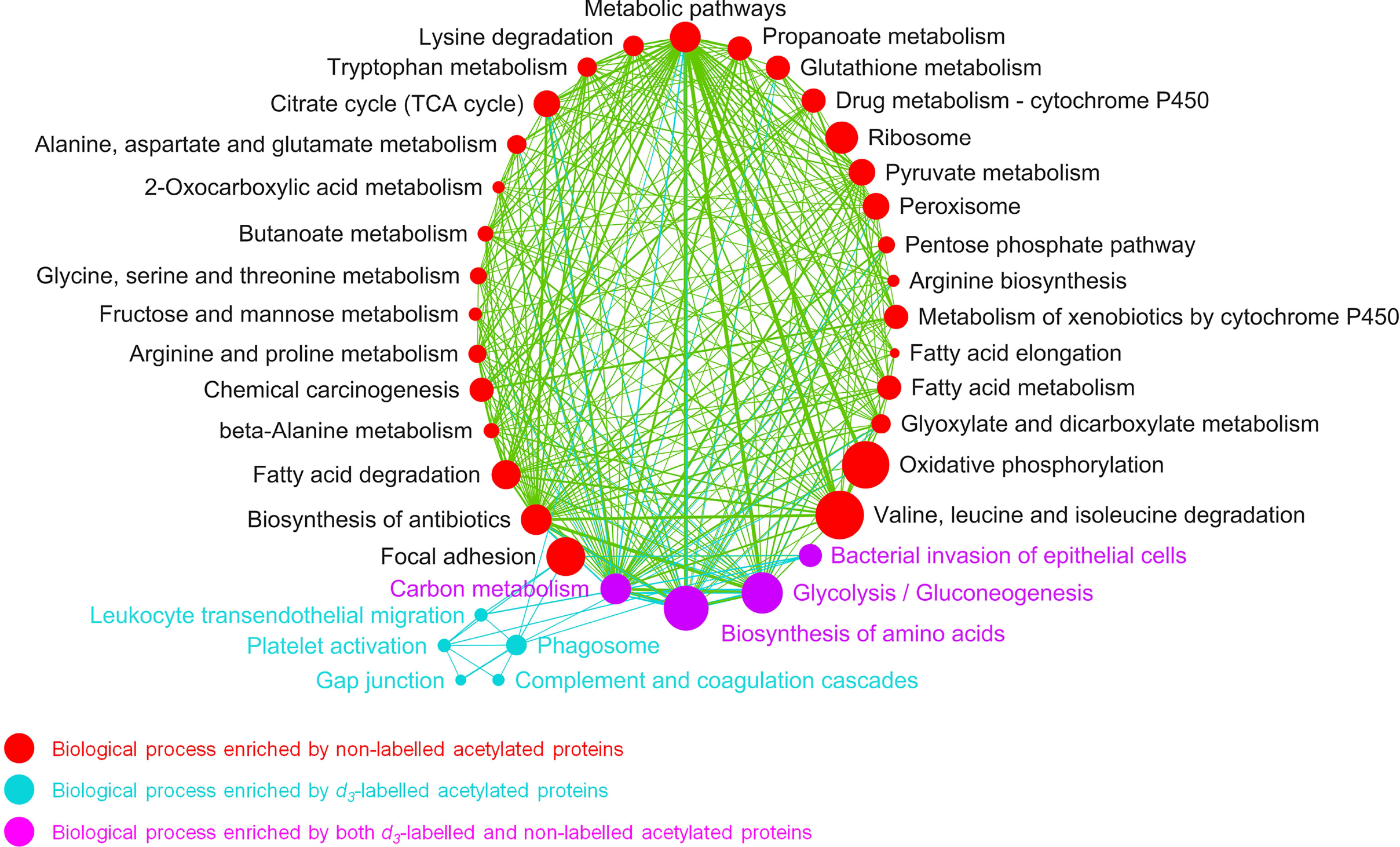
Biological processes enriched with proteins with ITM-acetylation in mice lung. Nodes stand for KEGG defined pathways (P<0.001 and FDR<0.001). Each node represents one biological process. The node size reflects the number of proteins in the pathway. Two nodes are connected if there were one or more proteins included in both nodes. Biological processes that are enriched by PTM-acetylated proteins (red), ITM-acetylated proteins (cyan), and proteins in both ITM-acetylome and PTM-acetylomes (plum) are highlighted in colors.

**Figure S5.**
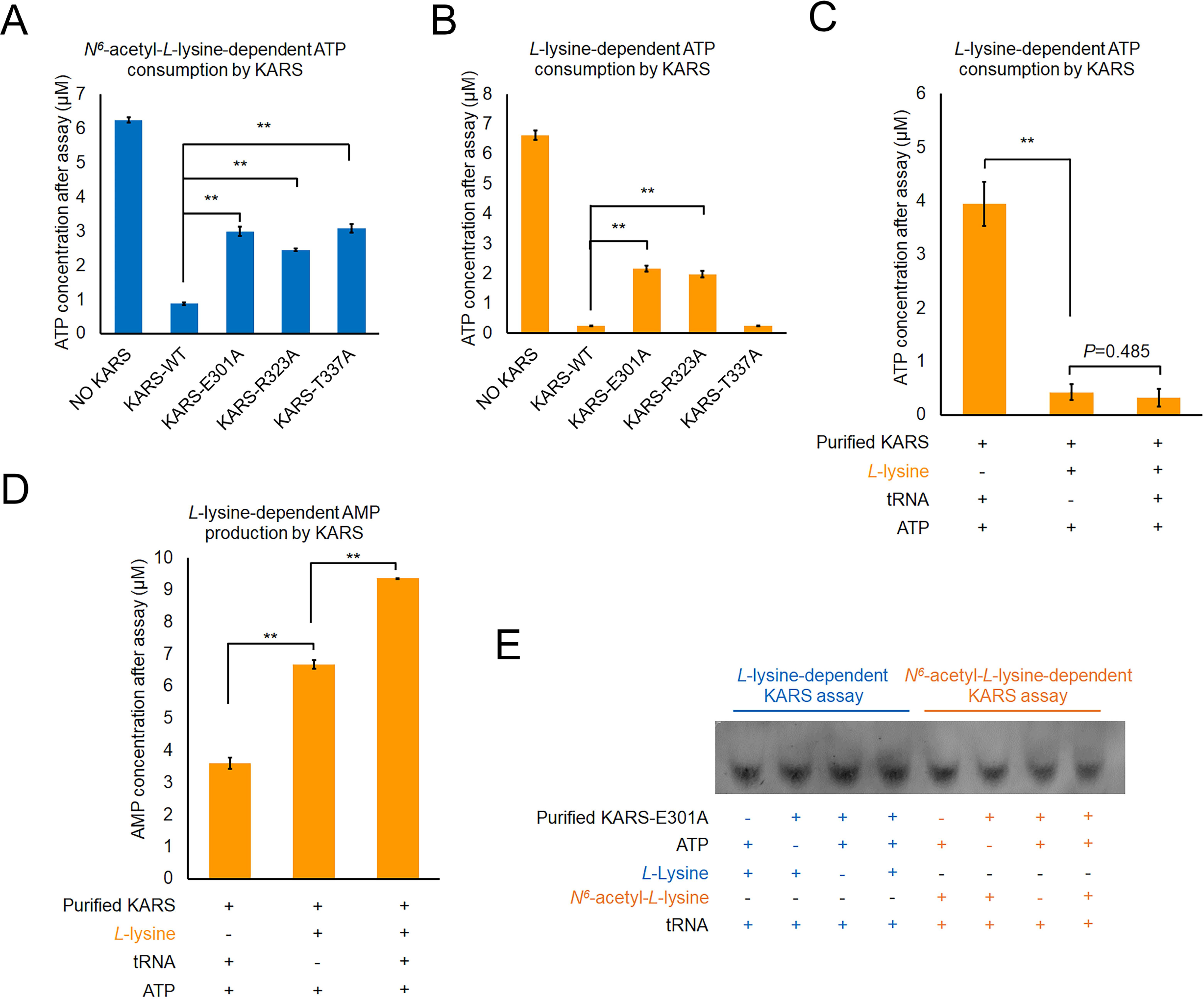
KARS can utilize *N*^6^-acetyl-*L*-lysine to produce *N*^6^-acetyl-*L*-lysyl-tRNA. (**A**) and (**B**) Identification of *N^6^*-acetyl-*L*-lysine and lysine interacting residues in the substrate binding pocket of KARS. Purified wild-type KARS and its mutants were incubated with ATP, tRNAs, and *N^6^*-acetyl-*L*-lysine (**A**) or *L*-lysine (**B**) or for 1 hour. The ATP consumption in the assays were measured and quantitatively analyzed. Two-sided *t*-test analyses were conducted. The data is presented as the means±s.d. of three independent experiments (n=3). \*\**P*<0.01. (C) and (**D**), KARS can catalyze ATP consumption and AMP production in the absence of tRNAs. Purified KARS was incubated with ATP and *L*-lysine for 1 hour. The ATP consumption (**C**) and AMP production (**D**) in the assays were measured and quantitatively analyzed. Two-sided *t*-test analyses were conducted. The data is presented as the means±s.d. of three independent experiments (n=3). \*\**P*<0.01. (**E**), KARS-E301A mutant losses the activity of catalyzing the formation of aminoacyl-tRNAs by using *L*-lysine and *N*^6^-acetyl-*L*-lysine. Northern blotting analysis of lysine cognate tRNAs by using fluorescence-labelled single strand DNA probe that specifically recognizes lysine cognate tRNAs. Representative images of triplicate experiments are shown.

**Figure S6.**
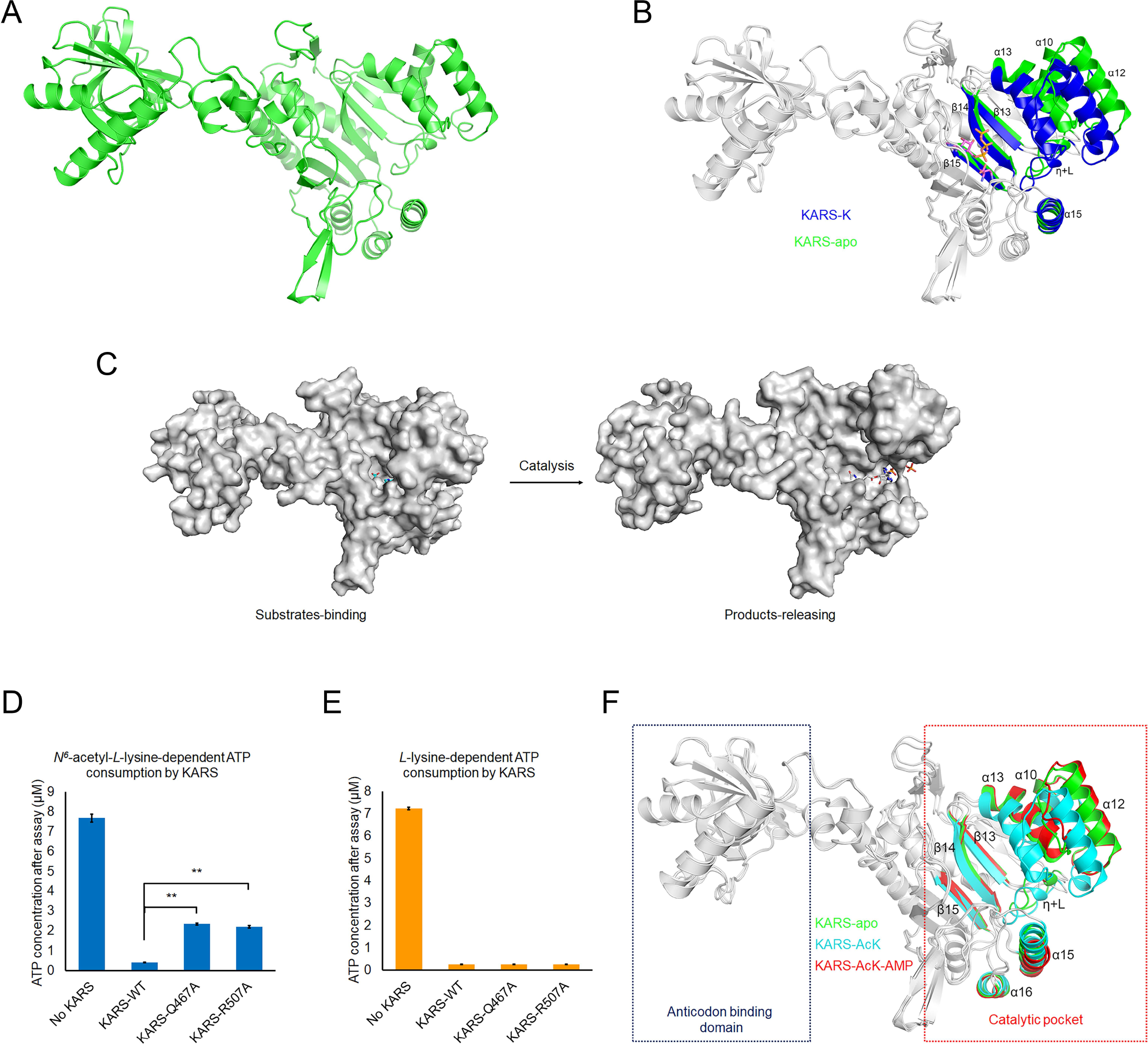
The molecular basis of KARS utilizing *N*^6^-acetyl-*L*-lysine as substrate. (**A**), Crystal structure of KARS-apo. Cartoon drawing of KARS-apo structure. (**B**), Structural difference between KARS in the states of standby and *L*-lysine-binding. Superimposition the structures of KARS-apo and KARS-K is performed. Only the regions showing structural difference between the KARS-apo (green) and KARS-K (blue) are colored and labelled, including the helixes of α-10, α-12, α-13, and α-15, the helix η and its associated loop (η+L), and the sheets of β-13, β-14, and β-15. *L*-Lysine and ATP are shown as sticks. (**C**), Surface drawing of the structures of KARS in the states of substrate-binding (KARS- AcK, left) and products-releasing (KARS-AcK-AMP, right). AcK, *N^6^*-acetyl-*L*-lysyl-AMP, and pyrophosphate are shown in sticks. (D) and (**E**), Gln467 and Arg507 are important to the *N^6^*-acetyl-*L*-lysine-dependent KARS activity. The Gln467 and Arg507 are mutated to Alanine, respectively. The in vitro KARS activity assays showed that Gln467 and Arg507 mutations significantly reduce *N^6^*-acetyl- *L*-lysine-dependent KARS activity (**D**) but have no influence on the lysine-dependent KARS activity (**E**). Each data is presented as the means±s.d. of three assays (n=3). Two-sided *t*-test analyses were conducted. \*\**P*<0.01. (**F**), Structure alignments of KARS-apo, KARS-AcK, and KARS-AcK-AMP. Superimposition the structures of KARS-apo, KARS-AcK, and KARS-AcK-AMP is performed. Only the regions showing structural difference between the KARS-apo (green), KARS-AcK (cyan), and KARS-AcK-AMP (red) are colored and labelled, including the helixes of α-10, α-12, α-13, α-15, and α-16, the helix η and its associated loop (η+L), and the sheets of β-13, β-14, and β-15. The compounds of *N^6^*-acetyl-*L*-Lysine, *N^6^*-acetyl-*L*- lysyl-AMP, and pyrophosphate are removed for better illustration of the protein structures.

**Figure S7.**
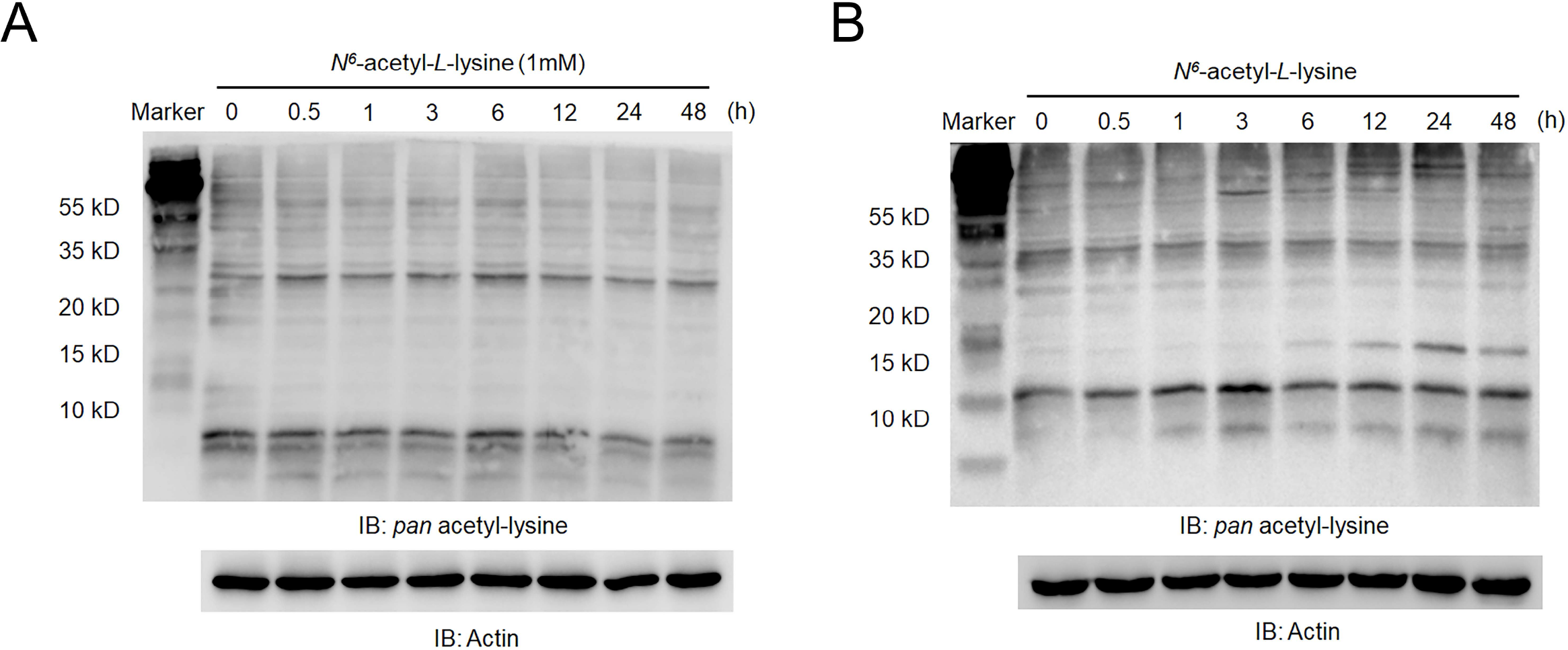
Deacetylases might surveil the influence of *N*^6^-acetyl-*L*-lysine on the acetylome in cells. (**A**), *N*^6^-acetyl-*L*-lysine treatment has no influence on the *pan*-acetylation of proteome in cells. The cultured LO2 cells were treated with 1 mM *N*^6^-acetyl-*L*-lysine for a series of time-points. The *pan*-acetylation of proteins in the treated cells were analyzed by immunoblotting assays with the indicated antibodies. Representative images of triplicate experiments are shown. (**B**), *N*^6^-acetyl-*L*-lysine treatment increases the *pan*-acetylation of proteins in cells treated with deacetylase inhibitor cocktail. The cultured LO2 cells were treated with 1 mM *N*^6^- acetyl-*L*-lysine for a series of time-points. Deacetylase inhibitor cocktail is added in the experiments to avoid the surveillance of deacetylases that might possibly remove the AcK treatment-induced protein acetylation in cells. The *pan*-acetylation of proteins in the treated cells were analyzed by immunoblotting assays with the indicated antibodies. Representative images of triplicate experiments are shown.

**Figure S8.**
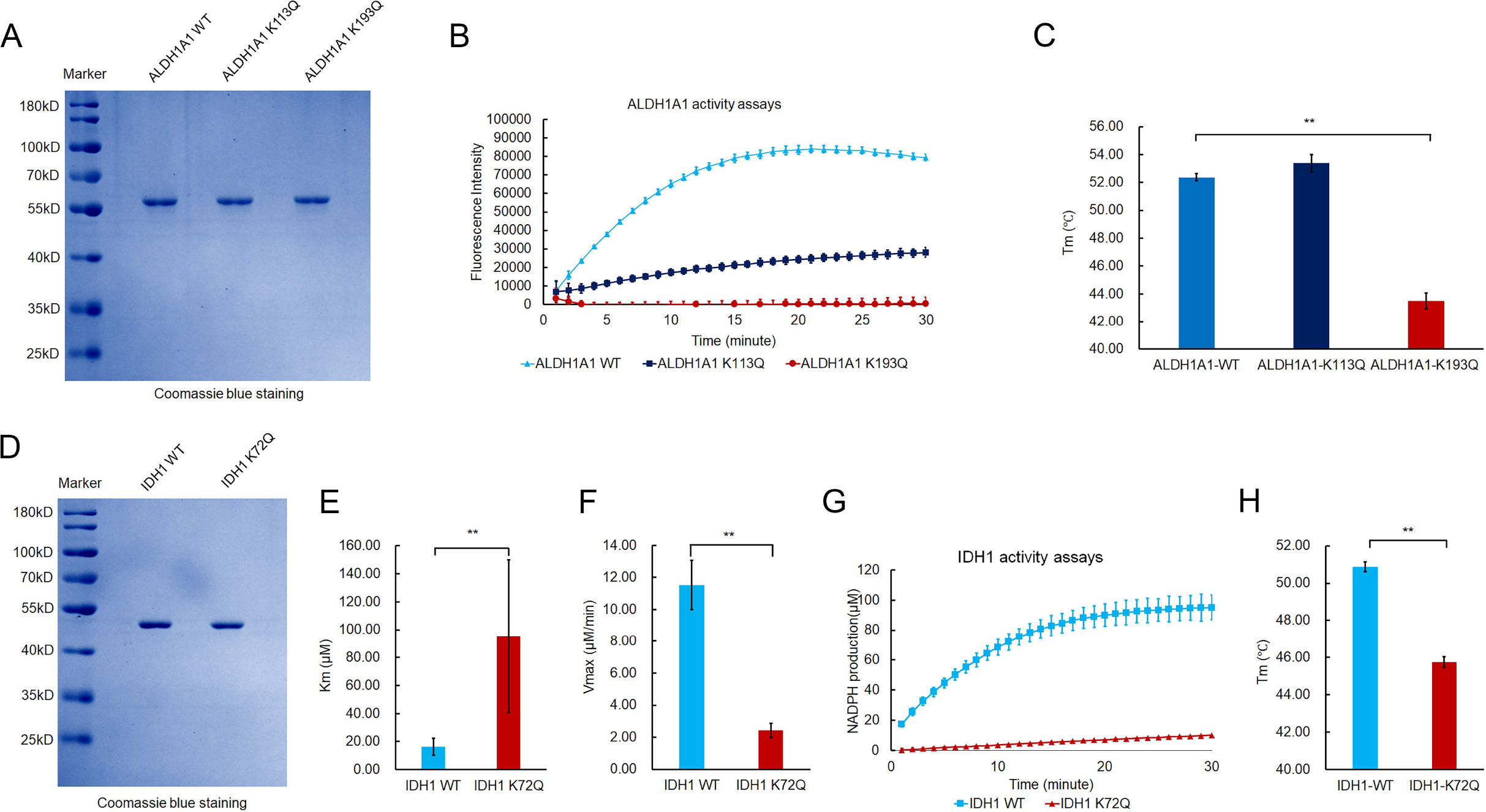
ITM can deposit acetylation in the PTM-cold regions of proteins. (**A**), Coomassie blue staining shows the purified protein of ALDH1A1wild-type and ALDH1A1 ITM-acetylation mimic mutants (ALDH1A1 K113Q and ALDH1A1 K193Q, respectively). (**B**), Acetylation mimic mutations of Lys113 and Lys193 reduce activity of ALDH1A1. The enzymatic activity assays of ALDH1A1 wild-type and mutants are performed by measuring the NADH production in each assay. Each data is presented as the means±s.d. of three independent assays (n=3). (**C**), Acetylation mimic mutation of Lys193 reduce the thermostability of ALDH1A1. The temperature of melting of ALDH1A1 wild-type and mutants are measured by thermofluor shift assays. Each data is presented as the means±s.d. of four independent assays (n=4). (**D**), Coomassie blue staining shows the purified protein of IDH1 wild-type and ITM- acetylation mimic mutant (IDH1 K72Q). (E) and (**F**), Acetylation mimic mutation of Lys72 reduces the NADP-binding ability and enzymatic activity of IDH1. The Michaelis Constant (Km) (**E**) and maximum velocity (Vmax) (**F**) of IDH1 utilizing NADP to produce NADPH were calculated through the steady kinetics analysis of IDH1. Each data is presented as the means±s.d. of three assays (n=3). Two-sided *t*-test analyses were conducted. \**P*<0.05, \*\**P*<0.01. (G) Acetylation mimic mutation of Lys72 reduces the activity of IDH1. The enzymatic activity assays of IDH1 wild-type and mutants are performed by measuring the NADPH production in each assay. Each data is presented as the means±s.d. of three independent assays (n=3). (H) Acetylation mimic mutation of Lys72 reduces the thermostability of IDH1. The temperatures of melting of IDH1 wild-type and IDH1 K72Q are measured by performing thermofluor shift assays. Each data is presented as the means±s.d. of four independent assays (n=4).

